# Proximity labeling reveals novel interactomes in live Drosophila tissue

**DOI:** 10.1101/542795

**Authors:** Katelynn M. Mannix, Rebecca M. Starble, Ronit S. Kaufman, Lynn Cooley

## Abstract

Gametogenesis is dependent on intercellular communication facilitated by stable intercellular bridges connecting developing germ cells. During Drosophila oogenesis, intercellular bridges (referred to as ring canals) have a dynamic actin cytoskeleton that drives their expansion to a diameter of 10*μ*m. While multiple proteins have been identified as components of ring canals (RCs), we lack a basic understanding of how RC proteins interact together to form and regulate the RC cytoskeleton. We optimized a procedure for proximity-dependent biotinylation in live tissue using the APEX enzyme to interrogate the RC interactome. APEX was fused to four different RC components (RC-APEX baits) and 55 unique high-confidence preys were identified. The RC-APEX baits produced almost entirely distinct interactomes that included both known RC proteins as well as uncharacterized proteins. The proximity ligation assay was used to validate close-proximity interactions between the RC-APEX baits and their respective preys. Further, an RNAi screen revealed functional roles for several high-confidence prey genes in RC biology. These findings highlight the utility of enzyme-catalyzed proximity labeling for protein interactome analysis in live tissue and expand our understanding of RC biology.

**Summary Statement:** Here, we optimized a procedure for enzyme-catalyzed proximity labeling in live Drosophila ovary tissue to interrogate the Drosophila ring canal protein interactome.

## Introduction

Ring canals (RCs) are intercellular bridges connecting developing germline cells during both male and female gametogenesis. Unlike other characterized RCs, Drosophila female germline ring canals accumulate an extensive F-actin cytoskeleton that drives their circumferential expansion during development from a ~1 *μ*m-diameter arrested cleavage furrow to an impressive 10 *μ*m-diameter tunnel. Genetic analyses have led to the identification of key proteins that are involved in ring canal structure (for a recent review, see Yamashita, 2018). Interestingly, proper ring canal development requires the ordered addition and/or activity of key proteins (Robinson, Cant, and Cooley, 1994; Cooley, 1998; Haglund, Nezis, and Stenmark, 2011), highlighting the dynamic and complex nature of the ring canal interactome. We also know that various ring canal proteins are regulated through post-translational modifications (PTMs) – with ubiquitination and phosphorylation being the most well-characterized PTM players at ring canals to date (Hudson, Mannix, Gerdes, et al., 2019; Morawe et al., 2011; Hamada-Kawaguchi, Nishida, and Yamamoto, 2015; Yamamoto et al., 2013; Kelso, Hudson, and Cooley, 2002; Kline, Curry, and Lewellyn, 2018).

While it is evident that many ring canal proteins are regulated in spatially-restricted and temporally-defined contexts, we still have an incomplete understanding of how ring canals are formed and regulated during development. Because of their insoluble nature and heterogeneity in size, classical biochemical purification and fractionation approaches that rely on abundant starting material in order to isolate and analyze ring canal proteins in native conditions are not practical. However, proximity labeling approaches have recently been developed (for reviews see Kim and Roux, 2016; Chen and Perrimon, 2017) in which an enzyme is fused to a protein or targeted to a sub-cellular compartment of interest where it can generate reactive ‘handle’ molecules or ‘tags’ (usually biotin-based) that covalently interact with proximal proteins in live cells. The resulting biotin-tagged proteins can then be purified by conventional methods and identified by mass spectrometry (MS). Importantly, these new techniques allow for the identification of proteomes and interactomes within the cell’s native environment. Additionally, the use of biotin tags makes the interrogation of insoluble cellular compartments possible since denaturing conditions can be used to solubilize, capture, and identify previously-inaccessible proteins (owing to the strength of biotin–streptavidin non-covalent binding; (K_d_≈10^−15^ M; Green, 1975).

The two most common classes of enzymes used for proximity labeling approaches are a bacterial biotin ligase (BirA) and peroxidase-based enzymes (horseradish peroxidase, HRP, and ascorbate peroxidase, APEX). BirA (Roux et al., 2012) and its newer, optimized variants (Kim, Jensen, et al., 2016; Branon et al., 2018) have been most commonly used for ‘BioID’ studies of protein-protein interactions, whereas HRP- and APEX-based applications have been used almost exclusively to identify subcellular organelle proteomes (for a more comprehensive review, see Kim and Roux, 2016). BioID uses membrane-permeable biotin for the substrate, but BirA’s relatively low enzymatic activity requires labeling reaction times of hours to days and an optimal temperature of 37°C, which is not practical for use *in vivo* in flies and worms. In contrast, APEX uses a less membrane-permeable substrate, biotin-phenol, but its high enzymatic activity allows for short labeling reaction times (seconds to minutes) at a wider range of temperatures. However, the membrane-impermeability of biotin-phenol has posed a significant technical challenge for use of APEX in live tissue.

Here, we report the application of APEX-mediated proximity labeling to the study of protein-protein interactions (PPIs) in the highly regulated but largely insoluble F-actin cytoskeleton of Drosophila female ring canals. We fused APEX to known ring canal proteins (‘RC–APEX baits’) to probe their interaction partners and identified high-confidence interactors (‘preys’) both unique to each RC–APEX bait or common to multiple RC–APEX baits. We used a proximity ligation assay and genetic analysis to validate the high-confidence preys. This work demonstrates that APEX-mediated biotinylation can be used to uncover specific PPIs within an intact cytoskeletal structure in living cells.

## Results

### Use of APEX to identify ring canal protein interactomes in live tissue

To interrogate ring canal protein interactomes, we made constructs in which APEX was fused to the ring canal proteins Pavarotti (Pav), HtsRC, Kelch, and a substrate-trapping Kelch construct, KREP (Figure 1A-B), and generated transgenic flies expressing these fusion proteins (RC–APEX baits): Pav::APEX, HtsRC::APEX, APEX::Kelch, and APEX::KREP (Figure 1B).

**Figure 1.**
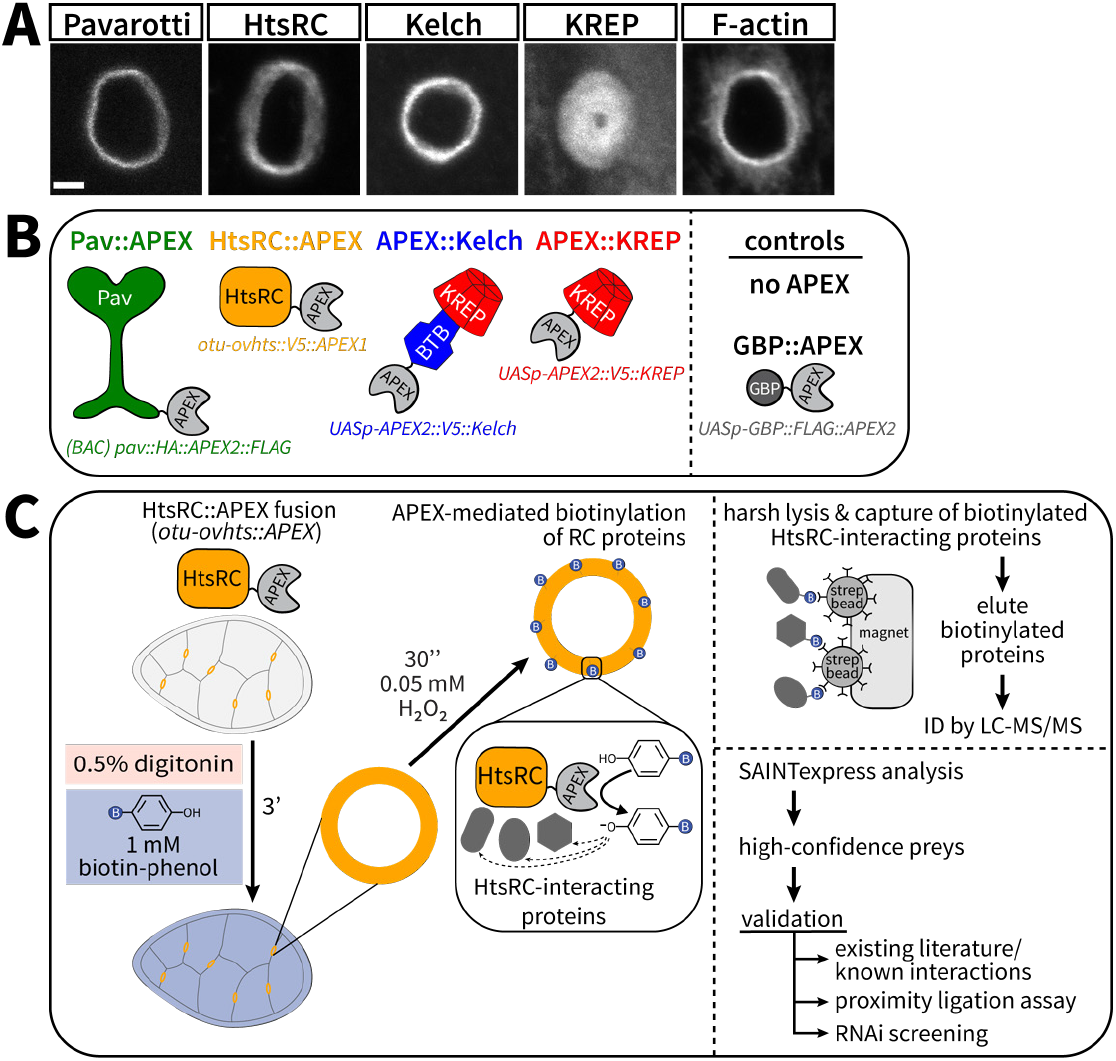
General workflow used to identify ring canal protein interactomes with APEX. (A) Fluorescence images showing localization of indicated proteins at ring canals. Pavarotti was visualized by GFP fluorescence of a (*BAC*) *pav*::GFP transgene. HtsRC and Kelch were visualized with anti-HtsRC and anti-Kelch antibodies, respectively. KREP was expressed as *UASp–APEX::V5::KREP* driven by *mat–GAL4* and visualized by anti-V5 immunofluorescence. F-actin was visualized by TRITC-phalloidin. Scale bar, 2 *μ*m. (B) Scheme of transgenic APEX fusion constructs generated for this study, with their corresponding genotypes listed below. Cartoons were made based on known structural models of Pav, Kelch, and KREP, with APEX positioned according to where it was fused. No structural data are available for HtsRC. (C) (Left) Example workflow of optimized APEX labeling in live ovary tissue with HtsRC::APEX construct. (Right) Overview of streptavidin purification and mass spectrometry analysis methods used to identify and validate high-confidence preys as novel protein interactors.

Pavarotti (Pav) is a kinesin-like protein known to be a component of all characterized germline ring canals as well as Drosophila somatic cell ring canals (Adams et al., 1998; Minestrini, Mathe, and Glover, 2002; Airoldi et al., 2011). Pav and its known binding partner Tumbleweed (Tum), a RacGAP, constitute the centralspindlin complex as a heterodimeric tetramer (Adams et al., 1998; Tao et al., 2016). HtsRC is a ring canal-specific protein required for recruiting the ring canal actin cytoskeleton (Petrella, Smith-Leiker, and Cooley, 2007; Hudson, Mannix, Gerdes, et al., 2019). Kelch is the substrate adaptor component of a Cullin3-RING ubiquitin ligase (CRL3) that targets HtsRC (Hudson and Cooley, 2010; Hudson, Mannix, and Cooley, 2015; Hudson, Mannix, Gerdes, et al., 2019). Since the KREP protein consists of only the substrate-targeting domain of Kelch, it cannot bind to Cullin3 and hence cannot ubiquitinate its substrate, HtsRC, which leads to a dominant-negative kelch-like ring canal phenotype (Hudson and Cooley, 2010; Figure 1A).

Pav localizes to the outer rim of ring canals (Minestrini, Mathe, and Glover, 2002; Ong and Tan, 2010), while HtsRC and Kelch are localize to the ring canal F-actin-rich inner rim (Robinson, Cant, and Cooley, 1994). Knowing that these ring canal proteins have discrete ‘sublocalizations’ within the ring canal as well as unique functions, our goal was to identify the local interactomes of each RC–APEX bait protein within the ring canal using APEX-mediated proximity biotinylation as the basis for subsequent proteomic analysis (see Figure 1C for general methods workflow and proteomic approach).

### Localization and expression of RC–APEX bait proteins

We matched the expression of the RC–APEX baits as closely as possible to their endogenous protein counterpart to minimize mislocalization that could lead to a high background and false-positive results (Hung et al., 2016; Varnaitė and MacNeill, 2016). For Pav::APEX, the APEX coding region was fused to the 3’-end of *pav* in the context of a bacterial artificial chromosome (BAC) encompassing the *pav* gene; previous C-terminal protein fusions to Pav did not perturb its function (Goshima and Vale, 2005). To generate HtsRC::APEX, the APEX coding region was fused to the 3’-end of the *ovhts* cDNA under control of the *otu* promoter, which closely matches the expression levels of the endogenous *ovhts* promoter (Petrella, Smith-Leiker, and Cooley, 2007). *APEX::kelch* and *APEX::KREP* were put under *UASp* control and expressed using *mat–GAL4* driver, which has been shown previously to be an appropriate GAL4 driver for *kelch* and *KREP* constructs (Hudson and Cooley, 2010; Hudson, Mannix, and Cooley, 2015; Hudson, Mannix, Gerdes, et al., 2019). We also made a control construct expressing unlocalized APEX by fusing GFP Binding Protein (GBP) (Rothbauer et al., 2008) to APEX under *UASp* control. Epitope tags (FLAG, HA, or V5) were included to allow visualization by immunofluorescence microscopy.

Immunofluorescence microscopy revealed that the RC–APEX bait constructs localized properly to ring canals (Figures 2A-D and S1A-D), and the GBP::APEX control was present throughout the cytoplasm of nurse cells (Figures 2E-E’ and S1E-E’). As expected, the APEX::KREP protein induced kelch-like ring canals (Figure 2D”, yellow boxed inset). Of note, Pav::APEX, HtsRC::APEX, and APEX::Kelch all rescued in their respective null mutant backgrounds (not shown), demonstrating that the RC–APEX fusion constructs were functional.

**Figure 2.**
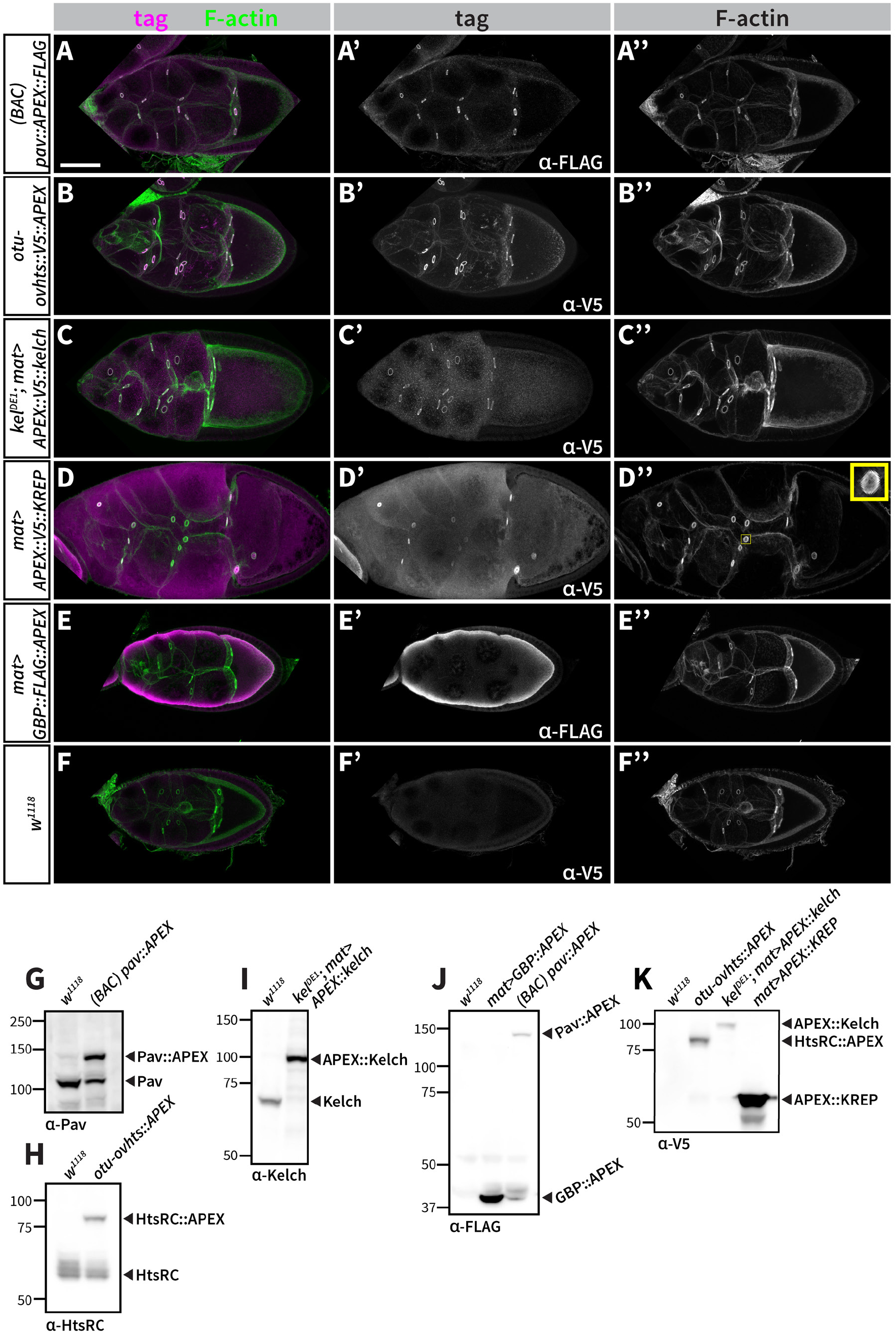
RC–APEX bait constructs localize to ring canals and are expressed at appropriate levels. (A - F) Stage 9 or 10 egg chambers expressing various RC–APEX baits, as indicated by genotypes listed on the left, were stained with TRITC-phalloidin (A”-F”) and epitope tags (A’-F’) to visualize and confirm proper RC–APEX bait construct localization to ring canals. Note the kelch-like ring canals induced by expression of APEX::KREP (D”, yellow boxed inset) and the cytoplasmic localization of the GBP::APEX control. Scale bar, 50 *μ*m. (G-K) Ovary lysates expressing indicated RC–APEX fusion baits were analyzed by western blotting to assess RC–APEX bait expression levels. Pav::APEX (G) and HtsRC::APEX (H) were expressed at levels closely matching endogenous Pav and HtsRC. (I) Levels of *APEX::Kelch*, expressed in a *kelch* mutant background, were comparable to endogenous Kelch. (J) *mat–GAL4-driven GBP::APEX* had higher expression than *(BAC) pav::APEX*, as visualized by FLAG immunoblotting. (K) *mat–GAL4-driven APEX::KREP* was the most abundantly expressed RC–APEX bait construct, consistent with previous studies showing high expression of similar KREP-derived constructs (Hudson and Cooley, 2010).

We directly compared expression of the RC–APEX constructs with their endogenous counterparts by western analysis (Figure 2G-K). Pav::APEX (Figure 2G) and HtsRC (Figure 2H) were expressed at comparable levels to endogenous Pav and HtsRC. APEX::Kelch protein expression was slightly higher than endogenous Kelch (Figure 2I), which was expected based on the moderately-high levels of expression produced by the *mat–GAL4* driver (Hudson, Mannix, and Cooley, 2015). We also compared expression of different RC–APEX constructs relative to each other using epitope tag antibodies. Detection of GBP::APEX and Pav::APEX with FLAG immunoblotting showed that GBP::APEX expression was significantly higher than Pav::APEX (Figure 2J), which was expected considering *mat-GAL4* was used to drive expression of *GBP::APEX*. Detection of HtsRC::APEX, APEX::Kelch, and APEX::KREP proteins with V5 antibodies showed that APEX::KREP was the most highly expressed of the three (Figure 2K), which is consistent with the high expression previously observed by other KREP-derived constructs (Hudson and Cooley, 2010). Overall, expression of the RC–APEX baits was comparable to endogenous levels.

### Biotinylation of ring canal proteins by RC–APEX baits and streptavidin purification

Most previous studies using APEX have been in cultured cells that are sufficiently permeable to the biotin-phenol APEX substrate after a 30 minute incubation period. The sole Drosophila APEX study (Chen, Hu, et al., 2015) used dissected Drosophila imaginal discs, salivary glands, and larval muscles – all of which were thin enough to allow biotin-phenol entry during the 30 minute incubation. The egg chamber, however, is within a muscle sheath and basement membrane, and the germline cells are surrounded by a layer of somatic follicle cells, thereby posing a challenge for the bulky biotin-phenol substrate (Rhee et al., 2013; Hung et al., 2016) to reach ring canals in the germline cells. Our initial experiments using the standard APEX labeling protocol developed by the Ting lab (Rhee et al., 2013; Hung et al., 2016) for use in cultured cells were unsuccessful. Suspecting a permeability issue, we pre-treated live egg chambers with a small amount of detergent to permit entry of the biotin-phenol substrate; this pre-treatment with detergent allowed for robust biotin labeling of ring canals (Figure S2).

After optimizing the parameters of the biotin-phenol labeling protocol (Figure 1C, left), we could achieve consistent and robust ring canal biotinylation with a three minute incubation in PBS (pH 8.0) with 0.5% digitonin and 1 mM biotin-phenol, followed by 30 seconds with 0.05 mM H_2_O_2_. We quenched the labeling reaction by addition of 10 mM NaN3, which inhibits the APEX enzyme. For visualization of biotinylation by immunofluorescence, egg chambers were fixed and stained with streptavidin-AF488 and TRITC-phalloidin (see Methods). Fluorescence microscopy revealed that the RC–APEX fusions specifically biotinylated ring canals in egg chambers through all stages of oogenesis, including stage 6 or 7 egg chambers (Figure 3) and stage 9 or 10 egg chambers (Figure S3). As expected, Pav::APEX also biotinylated follicle cell ring canals (Figure 3A’, yellow arrows), and GBP::APEX showed cytoplasmic biotin staining (Figure 3E’).

**Figure 3.**
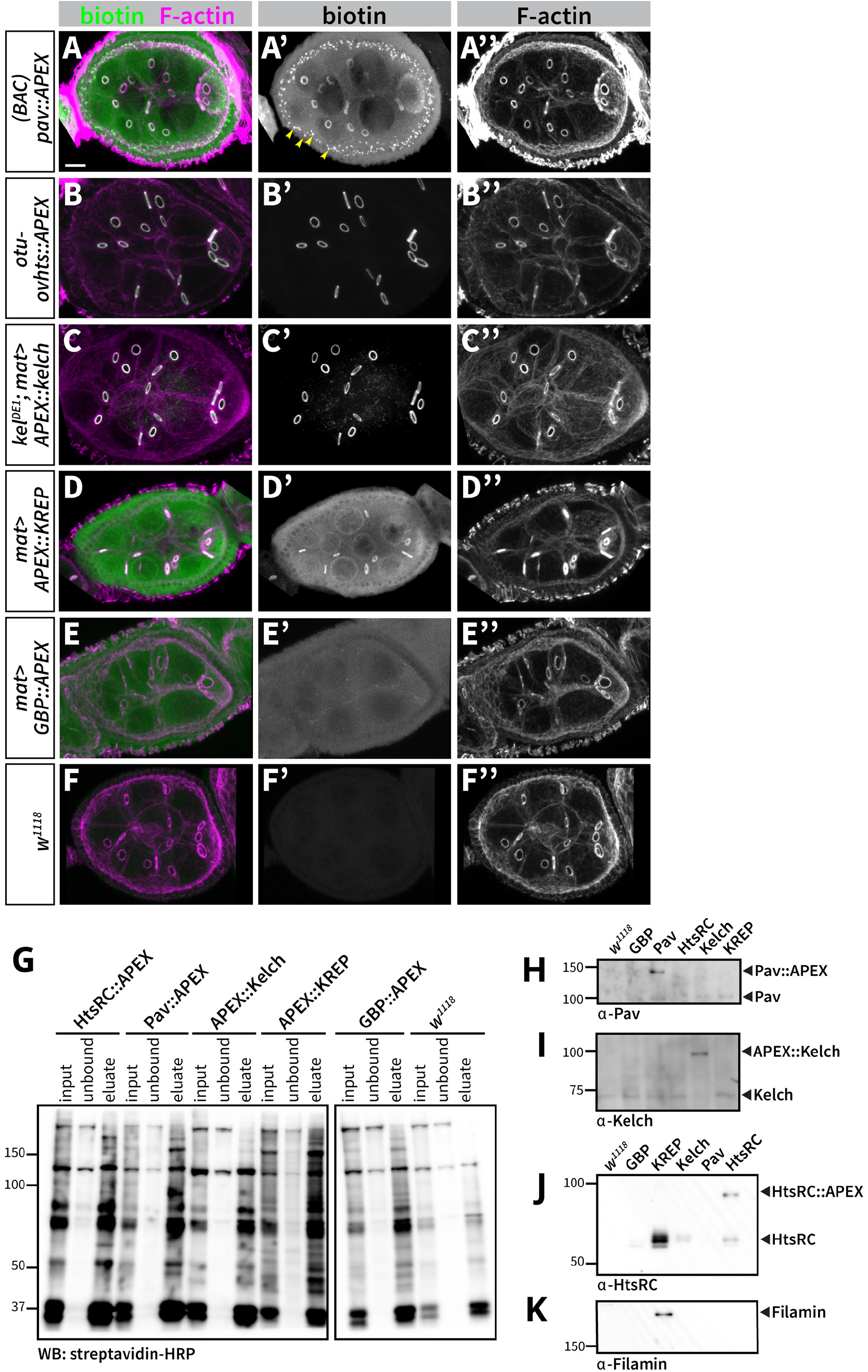
RC–APEX baits biotinylate distinct proteins in live egg chambers. (A-F) Ovaries expressing indicated RC–APEX baits were permeabilized and biotinylated, then egg chambers were fixed and stained with TRITC-phalloidin (A”-F”) and streptavidin-AF488 (A’-F’) to visualize F-actin and biotin. Representative images are shown of stage 6 or 7 egg chambers. Note the specific biotin signal at ring canals for the RC–APEX fusions (A’-D’) and the cytoplasmic biotin signal for unlocalized GBP::APEX (E’). As expected, the no-APEX control had no biotin signal (F’). Scale bar, 50 *μ*m. (G-K) APEX-labeled ovaries were lysed in denaturing conditions, biotinylated proteins were captured by magnetic streptavidin beads, and eluates were analyzed by western blotting. (G) Inputs, unbound fractions, and eluates were analyzed by western blotting with Streptavidin-HRP to confirm that biotinylated proteins were present and captured. (H-K) Western analysis was performed on the indicated APEX eluate samples (labeled on top of lanes) to check for the presence of known ring canal proteins. (H) Pav::APEX protein was abundant in the Pav::APEX eluate. (I) APEX::Kelch was enriched in the APEX::Kelch eluate, while a background endogenous Kelch protein band could be detected in all other samples. (J) HtsRC::APEX and endogenous HtsRC were present in the HtsRC::APEX sample, while endogenous HtsRC was highly-enriched in the APEX::KREP sample, consistent with previous work demonstrating KREP binds specifically to HtsRC (Hudson, Mannix, Gerdes, et al., 2019). (K) By Western analysis, Filamin was observed only in the APEX::KREP sample.

While treatment with detergent did not affect the structure, stability, or apparent composition of ring canals (Robinson and Cooley, 1997b) and thus was not expected to introduce any artifacts or alteration in the ring canal proteome, detergent treatment could pose a problem for studying cellular structures that are more labile. Since APEX was initially engineered and developed for use in electron microscopy (EM) studies in a procedure including fixation by 2% glutaraldehyde (Martell et al., 2012), we tested whether APEX remained active after formaldehyde fixation (Figure S4). We found that APEX retained activity even after a 10 minute fixation in 2% paraformaldehyde (Figure S4B, B”), suggesting APEX-mediated biotinylation will be broadly useful in fixed tissues. Nonetheless, to avoid potential fixation-induced cross-links that could obscure peptide identification, we proceeded using the protocol without fixative since the ring canals were stable even with detergent treatment.

In order to capture the biotinylated proteins using Streptavidin beads, we needed to make sure they were first solubilized after lysis and present in the soluble fraction after centrifugation and lysate clarification. Ring canals are known to be extremely stable structures that remain intact after lysis with non-ionic detergents such as TX-100 (Robinson and Cooley, 1997b; Greenbaum, Ma, and Matzuk, 2007). Therefore, we lysed the egg chambers in harsh, denaturing lysis buffer containing 8 M urea using a motorized spinning pestle in order to ensure that ring canal protein solubilization was as complete as possible. The clarified lysate was incubated with magnetic Streptavidin beads, and beads were subsequently washed with a 2 M Urea Wash Buffer. Streptavidin beads were eluted with SDS sample buffer supplemented with 10 mM biotin by boiling at 95° with shaking for 15 minutes. The eluate was analyzed by western analysis (see Methods for more specific details). Even in the denaturing 8 M Urea Lysis Buffer, the magnetic Streptavidin beads were able to capture the biotinylated proteins (Figure 3G), as shown by the depletion of the Streptavidin-HRP signal in the unbound fractions and the strong signal of biotinylated proteins in the eluates. Interestingly, the no-APEX control *w^1118^* sample also showed a smear of biotinylated proteins by western blot (Figure 3G), presumably due to the presence of endogenously biotinylated proteins or proteins that were biotinylated by other endogenous peroxidases. In any event, the signal corresponding to captured biotinylated proteins was stronger in the eluates for the APEX samples compared to the no-APEX *w^1118^* control, as would be expected by robust APEX-mediated protein biotinylation.

Before proceeding with mass spectrometry analysis, we performed additional western blots to check for the presence or absence of key ring canal proteins in the eluates (Figure 3H-K). If each RC–APEX bait biotinylates unique protein interactors, then known or suspected binding proteins should be present in each RC–APEX bait eluate and absent in the other eluate samples. The results were encouraging. When we tested for the presence of Pav in the eluates, we observed that only the Pav::APEX sample showed enrichment over controls for a Pav protein species (Figure 3H, Pav::APEX band), which is consistent with Pav being known to dimerize (Minestrini, Mathe, and Glover, 2002; Sommi et al., 2010; Tao et al., 2016). Similarly, only the APEX::Kelch sample showed enrichment for a Kelch protein species (Figure 3I, APEX::Kelch band), which is also consistent with Kelch dimerization, in this case via its BTB domain (Robinson and Cooley, 1997a; Hudson and Cooley, 2010; Hudson, Mannix, and Cooley, 2015). Note that APEX::Kelch was expressed in a *kelch* mutant background, which is why endogenous Kelch was absent from that lane. A faint band corresponding to endogenous Kelch was detectable in the other eluates, presumably from nonspecific binding to the beads. Our previous work showed that HtsRC is the CRL3^Kelch^ substrate (Hudson, Mannix, Gerdes, et al., 2019), so we looked for enrichment of HtsRC in the APEX::KREP eluate sample (Figure 3J). We found that HtsRC was significantly enriched in the APEX::KREP construct eluate, which is consistent with it being the CRL3^Kelch^ substrate. We also detected HtsRC and HtsRC::APEX in the HtsRC::APEX eluate, which suggests HtsRC is able to dimerize. Finally, we checked the eluate samples for the presence of Filamin, a key ring canal protein. Interestingly, Filamin was only detectable in the APEX::KREP sample (Figure 3K), suggesting that Filamin is also in close proximity to the Kelch substrate-targeting KREP domain. These results gave us proof-of-principle that the RC–APEX baits indeed biotinylate unique subsets of proteins – presumably their native interaction partners at the ring canal.

### Analysis of purified samples by mass spectrometry

To prepare samples for proteomic analysis, we subjected egg chambers from 100 flies (≈200 ovaries) of each genotype to our biotinylation labeling protocol. The labeled and quenched samples were frozen, lysed, purified with Streptavidin beads as described above and in Methods, and the Streptavidin beads were frozen at −80°C until subsequent MS analysis. Two biological replicates were used for each of the six samples (the five APEX constructs and one ‘no APEX’ control). We subjected our samples to LC–MS/MS by MS Bioworks using the ‘IP-works platform’, which entailed SDS-PAGE analysis of our purified bead samples followed by fractionation of each lane into 10 gel slices, in-gel trypsin digestion, LC–MS/MS using a ‘medium’ length liquid chromatography gradient, and Mascot database searching of the resultant MS/MS raw data within the UniProt Drosophila melanogaster proteome. The corresponding Mascot DAT files were parsed with Scaffold (Proteome Software) for subsequent data visualization and analysis. Spectral counts (SpC) were used as a semi-quantitative measure of relative protein abundance. See Methods for more detailed information on the mass spectrometry workflow and proteomic data analysis.

As a quality control check, we assessed the correlation of spectral counts of each identified protein across the biological replicates for each RC–APEX bait (Figure S5A). Linear regression analysis showed a strong correlation across replicates for each RC–APEX bait, as indicated by the large coefficient of determination (R^2^) values (between 0.90 and 0.92) (Figure S5A, top left of each graph). To assess how each RC–APEX bait sample differed in terms of its total protein identification coverage, we generated density plots of the natural log (ln) of the normalized spectral abundance factor (NSAF) values for all proteins identified by MS (Figure S5B). The density plots for all four constructs overlapped closely, indicating similar extent of coverage of spectral counts between the samples, and eliminating any concerns of instrumentation- and sampling-based artifacts or significant deviations in sample complexity or content.

### Identification and characterization of RC–APEX high-confidence preys

To establish a list of high-confidence preys for each RC–APEX bait, we used the Significance Analysis of INTeractome (SAINT) tool to analyze our proteomic data (Choi, Larsen, et al., 2011). SAINT, a common tool used in proteomic analysis (Choi, Liu, et al., 2012; Morris et al., 2014), is an algorithm that uses statistical modeling based on Bayesian probabalistic scoring to parse apart ‘true interactors’ from ‘false interactors’ in AP-MS datasets. We used a SAINTscore cut-off of ≥ 0.8 (corresponding to a 5% false discovery rate), which resulted in 55 high-confidence preys between the different RC–APEX baits (Figure 4A). Three proteins were deemed high-confidence preys for the Pav::APEX sample: Pav, Tum, and CG43333. HtsRC::APEX had five high-confidence preys: Msp300, Filamin, HtsRC, CG43333, and Cp7Fc. APEX::Kelch and APEX::KREP had nine and 43 high-confidence preys, respectively. Note that only a few preys (listed in the Venn diagram in Figure 4A) were shared between the different RC–APEX baits.

**Figure 4.**
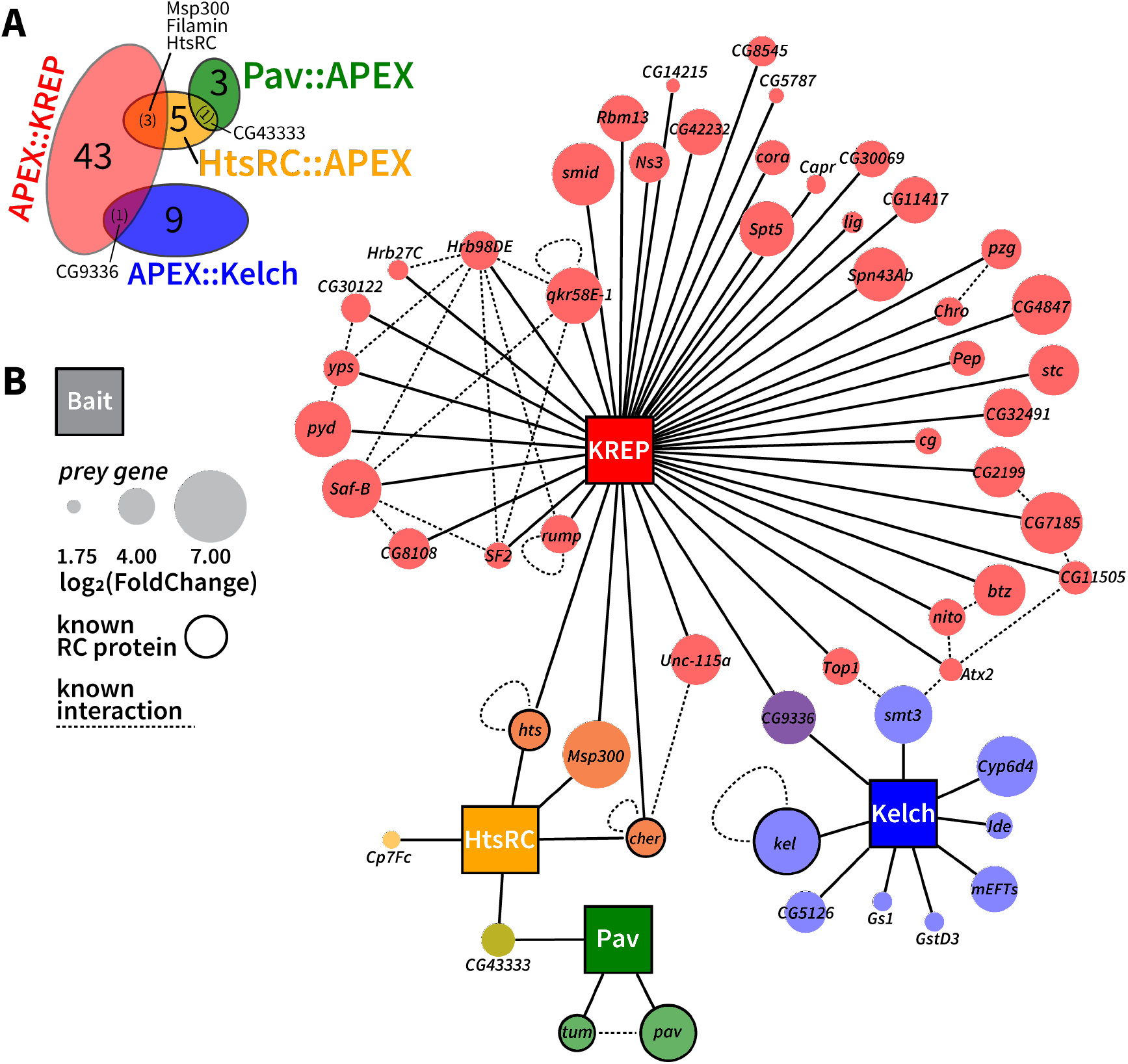
Proteomic analysis reveals RC–APEX baits have unique interactomes that include known ring canal proteins. (A) Venn diagram showing extent of overlap of the 55 high-confidence preys identified by the RC–APEX baits. High-confidence preys had a SAINTscore ≥ 0.8, using SAINTexpress analysis. (B) Network map generated in Cytoscape of the various RC–APEX baits, indicated by colored squares, and their respective prey genes. The size of the prey gene circle was dependent on its abundance in terms of spectral counts compared to control samples (log_2_(FoldChange)). Known ring canal proteins are denoted with a solid line outlining the prey gene circle. Dotted lines indicate previously-established interactions mined from the Molecular Interaction Search Tool (MIST) (Hu et al., 2018).

Next, we used Cytoscape to generate a network map of the RC–APEX baits and the genes for their respective high-confidence preys (Figure 4B). Of note, known ring canal proteins were identified as high-confidence preys (Figure 4B, solid line outlining prey gene circle). Consistent with our preliminary western analysis of RC–APEX bait eluates (Figure 3), Pav::APEX identified Pav; HtsRC::APEX and APEX::KREP identified HtsRC; APEX::Kelch identified Kelch; and APEX::KREP identified Filamin. Interestingly, Filamin was also identified by HtsRC::APEX even though it was not detectable by western analysis in the HtsRC::APEX eluate, which shows the sensitivity achieved by MS analysis. Pav::APEX also identified Tum, its known binding partner in the centralspindlin complex (Adams et al., 1998; Tao et al., 2016). To see if any of these prey genes were previously shown to interact with one another, we mined data from the Molecular Interaction Search Tool (MIST, an integrated resource for mining gene and protein interaction data; Hu et al., 2018), and imported the established known interactions into the Cytoscape network map (Figure 4B, dotted lines connecting prey genes).Taken together, these data support the idea that each RC–APEX bait identified unique interactomes that encompass previously-established interactors, new candidate interactors, and known ring canal proteins.

Of the 55 RC–APEX high-confidence prey genes identified, five were previously known to be involved in ring canal biology (Figure 4B, prey genes with solid outlines): *pav, tum, hts, cher*, and *kel*. That Pav::APEX identified Pav and Tum was reassuring, given that Pav dimerizes (Minestrini, Mathe, and Glover, 2002; Sommi et al., 2010) and also interacts with Tum (Adams et al., 1998; Tao et al., 2016). Kelch was the top prey for APEX::Kelch, which is consistent with Kelch dimerizing via its BTB domain (Robinson and Cooley, 1997a; Hudson and Cooley, 2010; Hudson, Mannix, and Cooley, 2015). And finally, HtsRC was identified by the CRL3^Kelch^ substrate-trapping APEX::KREP construct, which was also reassuring given that the CRL3^Kelch^ substrate is HtsRC (Hudson, Mannix, Gerdes, et al., 2019). Meanwhile, HtsRC::APEX identified HtsRC and Filamin as preys, and although HtsRC and Filamin are known ring canal proteins that genetically interact (Robinson, Smith-Leiker, et al., 1997; Sokol and Cooley, 1999), evidence of physical interaction between the two proteins had so far been elusive. Additionally, the fact that Filamin was a top prey for APEX::KREP was intriguing because it suggested that Filamin, HtsRC, and KREP may all be in a close proximity to one another at *kelch*-like ring canals.

To get a more global and quantitative visualization of the preys identified by each RC–APEX bait, we generated dot plots using the ProHits-viz tool (Knight et al., 2017) (Figure 5A-D). The color of each prey ‘dot’ is dependent on the Normalized Spectral Abundance Factor (NSAF) log_2_(FoldChange) for each bait compared to the control samples (see Figure 5 key). The size of each dot is dependent on its relative abundance between bait samples, meaning that the maximum-sized dot for each prey is allocated to the largest SpC value within that bait and the other dots are scaled proportionately. High-confidence preys within each bait have a yellow outline, corresponding to a SAINTscore equal to or greater than 0.8, while preys with a SAINTscore below 0.8 for that bait have a faint outline (see Figure 5 key). This type of visualization reveals how each RC–APEX bait identified unique sets of prey proteins. For example, APEX::Kelch identified nine high-confidence preys (shown in Figure 5C, third column of ‘dots’ all outlined yellow) and only one of those preys (CG9336) is shared with another RC–APEX bait (APEX::KREP) (yellow outline around CG9336 dot in fourth column for APEX::KREP).

**Figure 5.**
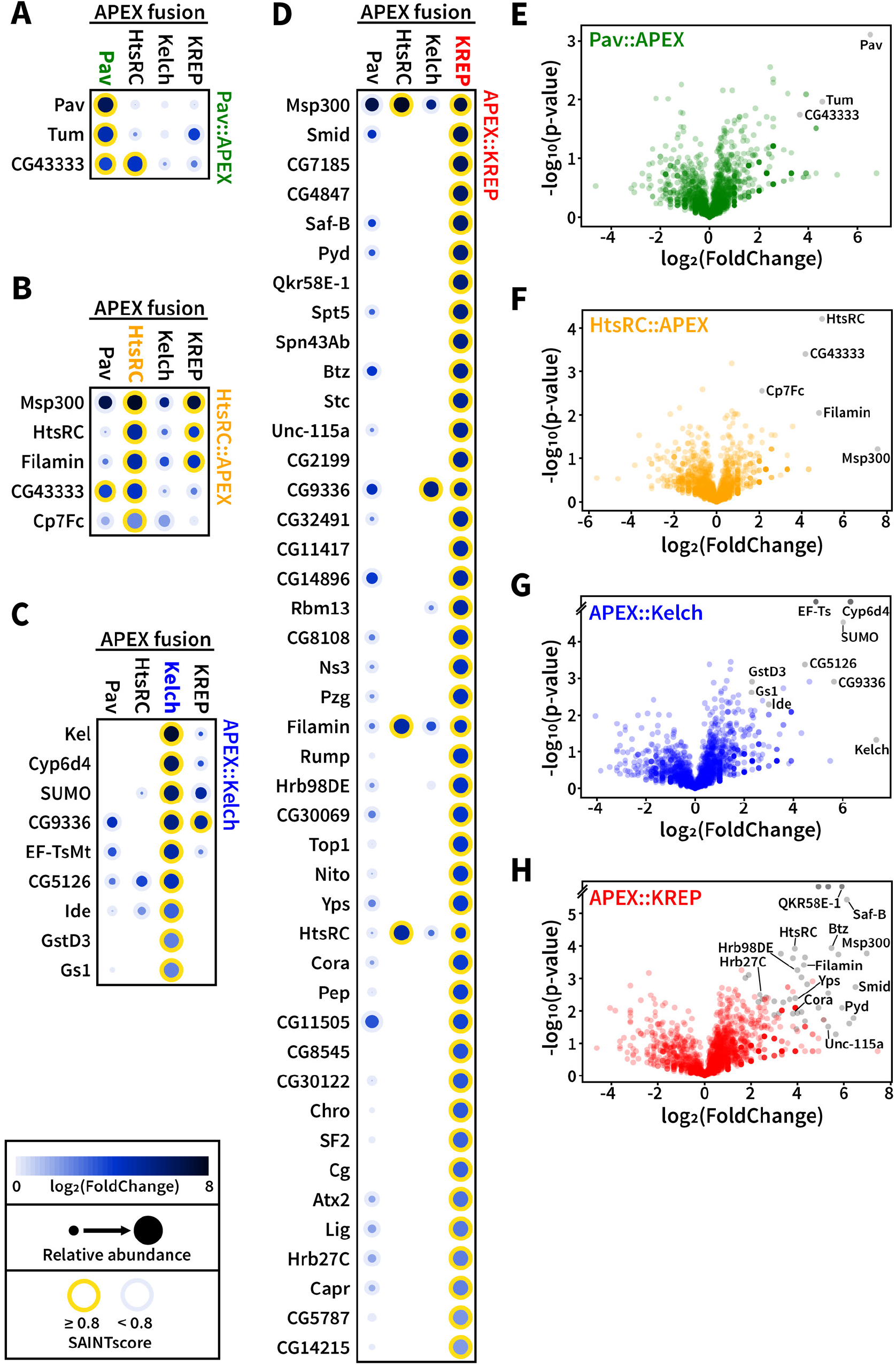
Proteomic analysis reveals RC–APEX baits identified unique prey proteins. (A-D) Dot plots of high-confidence preys identified by Pav::APEX (A), HtsRC::APEX (B), APEX::Kelch (C), and APEX::KREP (D). Dotplots were generated using ProHits-viz (Knight et al., 201), and high-confidence preys (SAINTscore ≥ 0.8) are outlined in yellow. Note that each RC–APEX bait identified mostly distinct high-confidence preys (compare columns). For example, in the APEX::Kelch column (C), only CG9336 was identified by another RC–APEX bait, APEX::KREP. (E-H) Volcano plots for each RC–APEX bait showing statistical significance (−log_10_(p-value)) and average relative abundance compared to controls (NSAF log_2_(FoldChange)) of identified proteins across replicate experiments. Notable high-confidence preys are listed and plotted as grey dots. Note for APEX::Kelch (G) and APEX::KREP (H), the y-axis is broken to allow room for the top-most row of protein preys that had an undefined p-value (e.g. Ef-Ts and Cyp6d4 in APEX::Kelch).

In order to visualize the quantitative proteomic data relative to its statistical significance, we generated volcano plots for each RC–APEX bait (Figure 5E-H). For each protein identified by MS, a p-value was calculated for its spectral counts within the biological replicates for each RC–APEX bait and control samples. For all identified proteins, the −log_10_(p-value) was plotted against the NSAF log_2_(FoldChange) compared to controls for each RC–APEX bait sample. In this type of scatter plot, proteins that showed a high fold-change compared to controls will appear on the right side of the plot, while proteins that are statistically significant compared to controls (in terms of SpC across replicates) will appear towards the top of the plot. Unsurprisingly, proteins in the top right of the plot generally corresponded to our SAINTexpress-generated high-confidence preys (Figure 5E-H, grey dots), since these should be robustly statistically significant with a large fold-change. For example, the five high-confidence preys identified by HtsRC::APEX are the sole occupants of the upper-right quadrant of the respective plot (Figure 5F, grey labeled dots).

### Validation of RC–APEX high-confidence preys by proximity ligation assay

Given that APEX-mediated biotinylation works in a proximity-dependent manner, the top preys should correspond to proteins that are in close proximity to the RC–APEX baits, and are therefore potential protein interactors or binding partners. Confirming physical interactions between these protein pairs in their native environments by conventional methods such as co-immunoprecipitation was essentially impossible due to the insoluble nature of the ring canal cytoskeleton. Therefore, we turned to the Duolink proximity ligation assay (PLA) to test if RC–APEX baits were in close proximity (i.e. <40 nm) to their respective prey proteins in egg chambers (see Methods for more information on PLA technology). We were able to test the five preys using available antibodies and GFP-tagged stocks. Initial PLA experiments using fixed whole egg chambers were unsuccessful. Suspecting a penetration issue, we performed the PLA in cryosectioned ovary tissue and succeeded in detecting PLA signals in 20 *μ*m ovary cryosections (see Methods for more details).

To test if Pav dimerization could be detected using the proximity ligation assay, we used egg chambers expressing the *pav::APEX::FLAG* BAC transgene. The presence of the FLAG epitope in the fusion protein allowed us to use FLAG antibodies generated in two species (mouse and rabbit) to target the same FLAG antigen. The PLA assay confirmed that Pav interacts with itself, as indicated by fluorescent foci throughout egg chambers and at ring canals (Figure 6A, top panel) compared to the negative control sample in which one of the antibodies was omitted (Figure 6A, bottom panel). We detected an interaction by the PLA between Pav and Tum using egg chambers expressing a *GFP::Tum* protein trap and the *pav::APEX::FLAG* BAC transgene. This genotype allowed us to use a mouse anti-FLAG antibody to target Pav::APEX::FLAG and a rabbit anti-GFP antibody to target GFP::Tum. Fluorescence microscopy showed a striking accumulation of PLA foci throughout the entire egg chamber and at the ring canals (Figure 6B, top panel), while the negative control was devoid of PLA signal (Figure 6B, bottom panel). Quantification of PLA foci showed that there were significantly more foci per egg chamber unit area in the {Pav}→Pav and {Pav}→Tum samples compared to their respective negative controls (Figure 6G, left graph). These results indicate that the proximity ligation assay can detect true protein-protein interactions at their native environment in tissue, and validates Pav and Tum as Pav::APEX preys.

We also tested the previously unknown potential interaction between HtsRC and Filamin in egg chambers expressing a *cher::GFP* protein trap using rabbit anti-GFP and mouse anti-HtsRC antibodies. Compared to the negative control, the experimental sample had abundant PLA foci throughout the egg chamber as well as a strong signal at ring canals (Figure 6C). Similarly, we tested potential oligomerization of HtsRC in egg chambers expressing the *otu-ovhts::V5::APEX* transgene, which allowed us to use mouse and rabbit antibodies targeting the same V5 antigen. We detected PLA foci throughout the egg chamber and at ring canals (Figure 6D). Quantification of PLA foci per egg chamber unit area showed that the {HtsRC}→Filamin and {HtsRC}→HtsRC samples had significantly more foci than their respective controls (Figure 6G, right graph). These results validate our proteomic results, and support the conclusion that HtsRC can dimerize or oligomerize as well as interact directly with Filamin, both in the cytosol and at ring canals.

**Figure 6.**
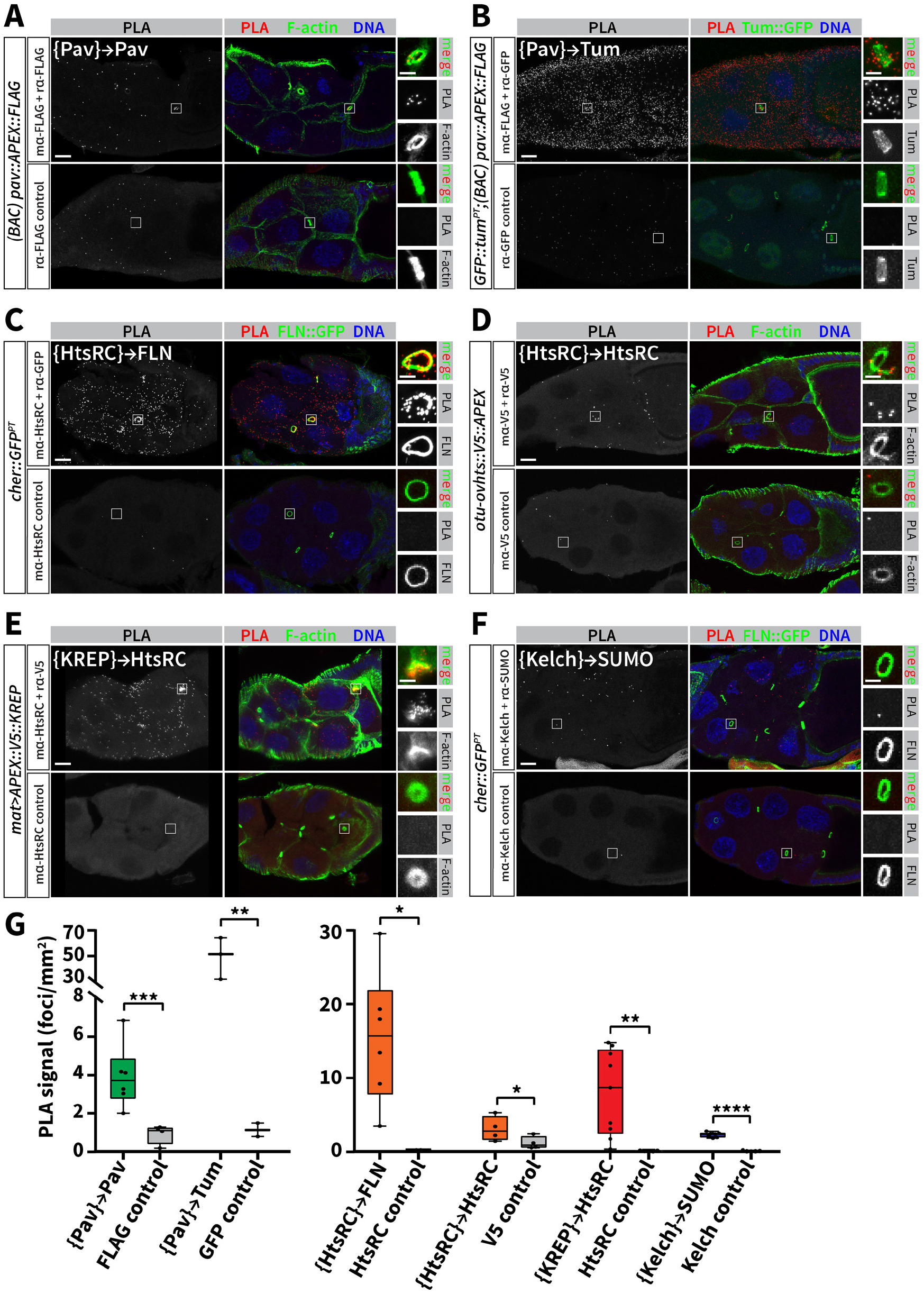
In situ proximity ligation assay confirms close-proximity interactions between RC–APEX bait proteins and respective preys. (A-F) Proximity ligation assay was performed on cryosectioned ovary tissue of the indicated genotypes to test for close-proximity interactions between different protein pairs, listed in white text in the top left of each panel (with brackets around the RC–APEX bait protein and an arrow pointed to the prey protein). A positive protein-protein interaction is indicated by the presence of PLA signal (left panels), which was visualized by fluorescent probes that bind to the PLA DNA product. Antibodies used for each reaction are indicated on the left, and the bottom panels served as negative controls since one antibody was omitted. Right boxed insets show ring canals, which were visualized by F-actin, GFP::Tum, or Filamin::GFP (FLN::GFP), as indicated in the panel labels. Note that {Pav}→Pav (A), {Pav}→Tum (B), {HtsRC}→Filamin (C), {HtsRC}→HtsRC (D), and {KREP}→HtsRC (E) all had positive PlA signal at ring canals, indicating a RC-localized protein-protein interaction between the respective protein pairs. The {Kelch}→SUMO (F) PLA signal was mostly dispersed throughout the cytoplasm. (G) Positive PLA foci in each egg chamber imaged were counted in FIJI using thresholding and particle analysis, and graphed as a function of egg chamber unit area. Student’s t-test was used to compare number of PLA foci per unit area in experimental samples vs. controls. *, p ≤ 0.1; **, p ≤ 0.05; ***, p ≤ 0.001; ****, p ≤ 0.0001. Scale bars, 20 *μ*m (main panel) and 5 *μ*m (insets).

HtsRC was a high-confidence prey for the substrate-trapping APEX::KREP construct, which was expected based on our work showing that HtsRC is the CRL3^kelch^ substrate (Hudson, Mannix, Gerdes, et al., 2019). We validated this result in egg chambers expressing *APEX::V5::KREP* using mouse anti-HtsRC antibody to target HtsRC and a rabbit anti-V5 antibody to target APEX::V5::KREP. As expected, strong PLA signal throughout the cytoplasm and at ring canals (Figure 6E) with essentially no foci in th enegative control (Figure 6E, bottom panel; quantified in Figure 6G, right). These results provide proof-of-principle that APEX fusion constructs can be used to identify substrates of E3 ubiquitin ligases in live tissue.

Finally, we tested an unexpected interaction between SUMO (encoded by the *smt3* gene) and Kelch using a Drosophila SUMO rabbit antibody (Abgent) and a mouse anti-Kelch antibody. We performed a PLA using a mouse anti-Kelch antibody and a rabbit anti-SUMO antibody. Excitingly, we observed PLA foci throughout the cytoplasm of egg chambers in the experimental sample but not the negative control (Figure 6F, compare top and bottom panels; quantification in Figure 6G, right). The results suggest that Kelch is in close proximity to the SUMO protein in the cytoplasm of egg chambers, but not at ring canals. Future experiments will be needed to test if Kelch is covalently SUMOylated or interacting with SUMO, perhaps through its predicted SUMO interaction motif (Zhao et al., 2014).

As an additional control, we used the PLA to test interactions between RC–APEX bait proteins and proteins that were *not* identified as preys. Specifically, we tested for interactions between Pav and Kelch (Figure S6A), Kelch and Filamin (Figure S6B), Pav and HtsRC (Figure S6C), and HtsRC and Kelch (Figure S6D). All interactions tested were negative as indicated by the absence of PLA foci, and quantification of the foci revealed that there was no significant enrichment of PLA foci in the experimental samples compared to their respective negative controls (Figure S6E). These results were all consistent with our RC–APEX bait and prey protein pairs and support our hypothesis that the PLA was faithfully recapitulating real protein-protein interactions.

### Identification of novel ring canal regulators through an RNAi screen of RC–APEX prey genes

To test genetically whether the RC–APEX high-confidence preys were involved in ring canal biology, we performed an RNAi screen using stocks from the Transgenic RNAi Project (TRiP). In total, we screened 33 prey gene RNAi lines using the *mat-GAL4* driver to drive expression of the shRNAs specifically in the female germline. Of the 33 genes screened, 19 had a noticeable phenotype; eight phenotypes were ring canal-specific and 11 had general oogenesis defects (Figure 7A). Six out of the eight genes that had ring canal-specific phenotypes (*btz, cg, Pep, pyd, SF2*, and *smid*) had significantly larger ring canals when their gene expression was knocked down by RNAi (Figure 7B-H; ring canal diameter quantified in Figure 7J). RNAi-mediated knockdown of *Pep* also led to abnormal F-actin structures as well as deformed and collapsed ring canals (represented by red, yellow, and blue arrows in Figure 7E’, respectively). Knockdown of *pyd* also resulted in deformed ring canals (yellow arrows in Figure 7F’), while *smid* knockdown revealed abnormal excess cortical F-actin surrounding ring canals in stage 10 egg chambers (red arrows in Figure 7H). *Spt5* knockdown led to egg chamber degeneration in mid-to-late staged egg chambers (Figure 7I) and collapsed ring canals (Figure 7I’, blue arrows). Finally, RNAi-mediated knockdown of *smt3* (the Drosophila SUMO gene) led to egg chamber arrest as well as deformed and collapsed ring canals (Figure 7K, yellow and blue arrows). To examine a possible role for SUMO in ring canal biology, we stained egg chambers with anti-SUMO antibody and found occasional ring canal localization (Figure 7L, boxed insets).

**Figure 7.**
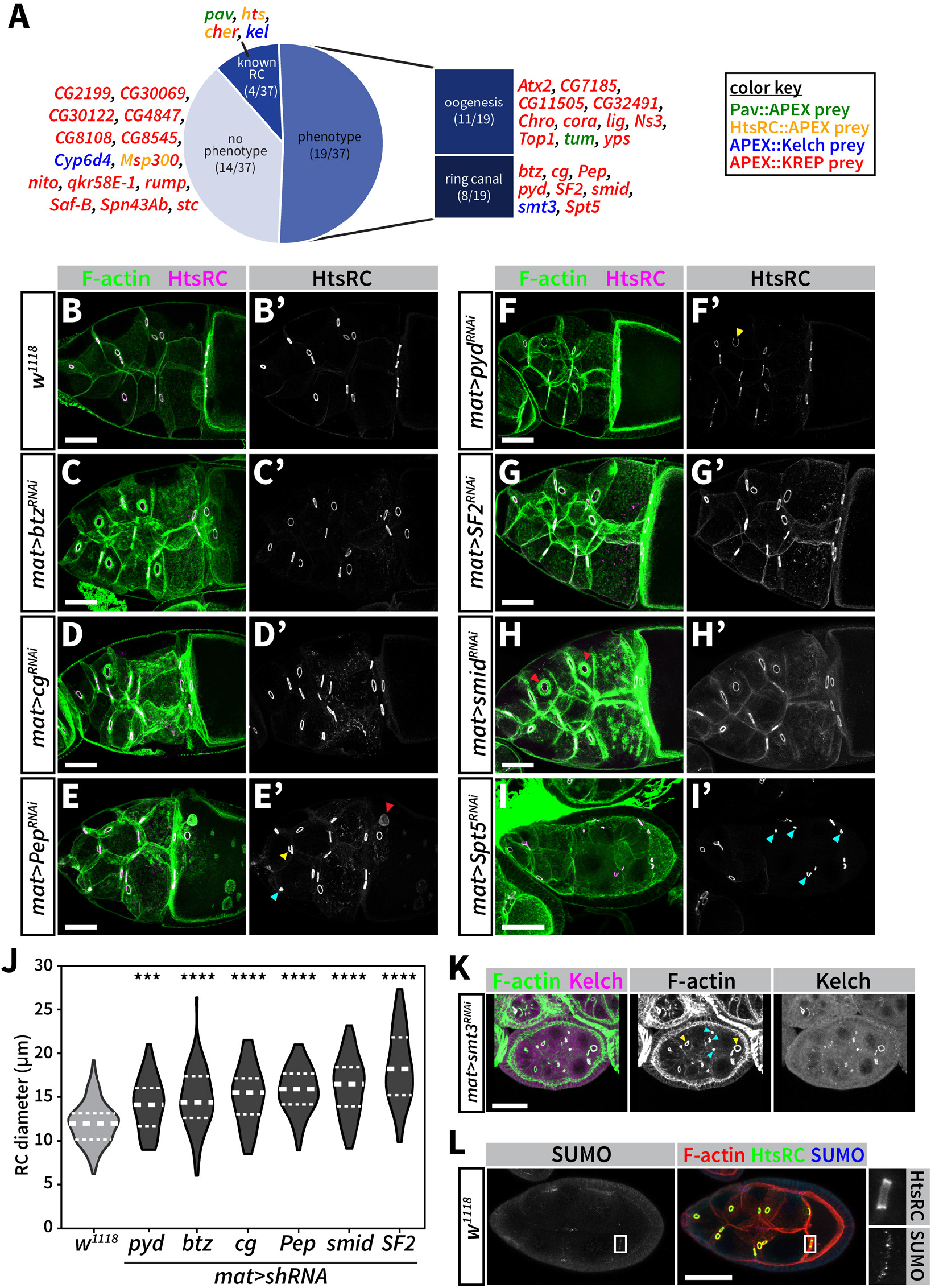
RNAi screen reveals prey genes are involved in ring canal biology. (A) Summary of results from RNAi screening of 55 high-confidence preys (with each prey color-coded to indicate its respective RC–APEX bait(s)). Of the 33 prey genes screened, 19 had a phenotype, eight of which were ring canal-specific. *pav, hts, cher*, and *kel* were not screened since their roles as ring canal proteins are well documented. (B-I) Egg chambers of indicated genotypes were stained with TRITC-phalloidin and HtsRC (B’-I’) to reveal ring canal size and morphology. Red arrows indicate abnormal F-actin structures. Yellow and blue arrows indicate deformed and collapsed ring canals, respectively. Scale bars, 50 *μ*m. (J) Ring canal diameters of the indicated genotypes were measured in FIJI using HtsRC staining as a ring canal marker, and violin plots were used to represent the data. Thick dashed white line shows median RC diameter, and thinner upper and lower white lines denote the upper and lower quartiles. One-way ANOVA was used to compare the mean RC diameter between the RNAi lines and *w^1118^* control. ***, p ≤ 0.0001; ****, p ≤ 0.00001. (K) RNAi-mediated knockdown of *smt3* leads to collapsed and deformed ring canals, marked by blue and yellow arrows, respectively. Scale bar, 25 *μ*m. (L) Some ring canals are positive for SUMO staining (inset). Scale bar, 20 *μ*m.

Interestingly, none of the eight genes with ring canal-specific phenotypes were known previously to have functions related to ring canals. We were particularly surprised to see that five of these genes (*btz, cg, Pep, SF2*, and *Spt5*) had established functions involving DNA and/or RNA binding, processing, or regulation. In summary, this preliminary RNAi screen revealed that prey genes were involved in ring canal biology, which provides additional support that the RC–APEX baits identified physiologically-relevant protein interactors. Future experiments will be necessary to gain mechanistic insight into how these genes are involved in ring canal biology.

## Discussion

### APEX-mediated proximity labeling can be used to identify protein-protein interactions in live tissue

Since its introduction in 2013, APEX-mediated localized biotinylation has been used successfully in a number of studies to identify organellar or cellular proteomes, with most of the work done in cultured cells. Three studies reported APEX-mediated labeling in live tissue. Two of these studies were performed in *C. elegans* in a mutant background that compromised cuticle integrity, which allowed for successful delivery of the biotin-phenol substrate into the worm tissue (Reinke, Balla, et al., 2017; Reinke, Mak, et al., 2017). The third labeled the mitochondrial proteome in dissected Drosophila imaginal discs, salivary glands, and larval muscles (Chen, Hu, et al., 2015). These tissues are relatively thin, which presumably permits sufficient entry of biotin-phenol into the cells without notable intervention. Here, we adapted the use of APEX-mediated proximity labeling to thicker tissue by using digitonin as a means to sufficiently permeabilize cell membranes and allow biotin-phenol substrate entry. Our approach could be adapted to other organisms and tissues, live or formaldehyde-fixed, to enable a more widespread use of APEX *in vivo*.

Of note, the Ting lab recently engineered more catalytically-active mutants of the BirA enzyme used in BioID approaches, referred to as TurboID and miniTurbo (Branon et al., 2018). Essentially, these mutants combine the catalytic efficiency of APEX (i.e. a 10 minute labeling reaction time) with the ease-of-use of BioID (i.e. straightforward biotin delivery to cells or tissues of interest). However, when TurboID was used in flies and worms (Branon et al., 2018), biotin supplementation took hours or days, which limits its potential for capturing more dynamic processes that require fine temporal control and resolution of the labeling reaction. In these cases, our APEX protocol could be used with dissected tissue to achieve robust biotinylation on a faster (seconds to minutes) timescale. Further adaptation and optimization of these proximity labeling approaches will undoubtedly open the door to proteomic experiments in living organisms.

### RC–APEX baits identified known binding partners as well as potential new interactors

We set out to test the feasibility of using APEX *in vivo* to identify protein-protein interactions as opposed to whole-organelle or whole-cell proteomes. While several RC proteins have been characterized, a full understanding of F-actin regulation in RCs requires much more complete information about PPIs in the structure. Because biotinylation of nearby proteins is dependent on their proximity to the APEX enzyme, we envisioned that the RC–APEX bait proteins could identify new binding partners. Our results were very promising. We were especially excited that the two top preys of Pav::APEX were Pav and Tum since identifying the Centralspindlin complex (Tao et al., 2016) provided a powerful positive control and a compelling proof-of-principle that the RC–APEX baits were indeed capturing real protein interactors of the respective ring canal bait proteins. Our results also indicated that APEX fused to different domains within protein can identify domain-specific interactors. APEX fused to the Kelch N-terminus, which is the site of Kelch homodimerization through its BTB domain (Errington et al., 2012; Canning et al., 2013; Ji and Prive, 2013) labeled distinct proteins compared to APEX::KREP, which contains just the substrate-binding domain of Kelch. Kelch itself was a top prey for APEX::Kelch while HtsRC was a top prey for APEX::KREP. Surprisingly, APEX::Kelch did not identify Cullin3, a known binding partner of the Kelch BTB domain (Xu et al., 2003; Errington et al., 2012; Canning et al., 2013; Ji and Prive, 2013), as a top prey. This might be due to the transient nature of the Kelch–Cullin3 interaction or the relatively low abundance of Cullin3 present at the ring canals compared to its strong cytoplasmic localization (Hudson and Cooley, 2010; Hudson, Mannix, and Cooley, 2015).

Identification of E3 ubiquitin ligase substrates is challenging because substrates are short-lived and their interactions with the ligase are transient. Those challenges, coupled with the insolubility of the ring canal actin cytoskeleton, made identifying the CRL3^Kelch^ ring canal substrate particularly hard. We recently identified HtsRC as the substrate of CRL3^Kelch^ with other methods (Hudson, Mannix, Gerdes, et al., 2019); however, our work presented here suggests APEX is a valuable new tool for identifying E3 ubiquitin ligase substrates, particularly in previously-inaccessible contexts like in live tissue or within insoluble cellular compartments such as the cytoskeleton. In fact, ours is the first study of its kind to use APEX *in vivo* to identify an E3 ubiquitin ligase substrate. The advent of substrate-trapping constructs that can be fused to APEX and expressed in model organisms could lead to future breakthroughs for substrate identification in tissue-specific and developmental contexts, which will help move the E3 ubiquitin ligase field into physiologically-relevant models.

In addition to the well-established protein-protein interactions captured by the RC–APEX baits, we also identified many unexpected proteins as high-confidence preys. Interestingly, many APEX::KREP prey genes are involved in RNA binding or processing as well as ribonucleoprotein complexs. RNPs are transported along microtubules, through ring canals, from the nurse cells into the oocyte (Mische et al., 2007). Given that expression of KREP leads to a dominant-negative phenotype that induces *kelch*-like ring canals, it is possible that the RNP complexes are being caught or trapped in the *kelch*-like ring canal cytoskeleton during their transport, which could have allowed their biotinylation and subsequent identification by APEX::KREP. However, RNAi-mediated knockdown of genes for several of these RNP-associated APEX::KREP prey genes led to ring canal-specific phenotypes, suggesting a yet-unappreciated functional role for mRNA regulation and RNP complexes in ring canal biology. Since these proteins were identified by APEX::KREP, whose expression essentially phenocopies a *kelch* null phenotype, the emergence of these RNA-regulating proteins in this background could reveal a novel and unexpected role for RNA and translational regulation that contributes to the *kelch*-like phenotype.

### Proximity labeling revealed new insight into ring canal substructure

Ring canals are massive structures that undergo dramatic and dynamic structural and possibly compositional changes during egg chamber development. Given the complex nature of the ring canal cytoskeleton (e.g. F-actin-dependent membrane and ring canal expansion, actin recruitment, F-actin crosslinking and bundling, cytoskeleton disassembly in the ring canal lumen), it would make sense that different sub-complexes exist in ring canals to carry out these processes. Evidence already exists for distinct domains within the ring canal. HtsRC, Kelch and F-actin localize to the inner rim of ring canals (Robinson, Cant, and Cooley, 1994), Pav localizes to the outer rim (Minestrini, Mathe, and Glover, 2002; Ong and Tan, 2010), and Filamin is present throughout the entire ring canal cytoskeleton. However, a more finely-resolved picture of ring canal protein complex localization is emerging from our localized biotinlyation results.

Our data showed that different RC–APEX bait constructs identified unique subsets of ring canal proteins suggesting there are sub-complexes within the ring canal that orchestrate mechanical functions such as nucleating actin filament assembly at the membrane to expand the ring canal diameter or controlling HtsRC levels to maintain an open ring canal lumen. Of particular interest is the finding that HtsRC::APEX identified Filamin and HtsRC as top preys. Given that localization of HtsRC to RCs is genetically dependent on the *cher* gene (Robinson, Smith-Leiker, et al., 1997; Sokol and Cooley, 1999), we can now speculate that Filamin directly recruits HtsRC to the ring canal where we know HtsRC functions to recruit the abundant F-actin cytoskeleton (Hudson, Mannix, Gerdes, et al., 2019). This also suggests that HtsRC has the ability to dimerize or oligomerize, possibly through its C-terminal coiled-coil domain (predicted using Phyre2; Kelley et al., 2015). It will be interesting to test if oligomerization status of HtsRC differs based on its localization in the cytoplasm or at ring canals and whether this affects its actin-regulating activity.

It is also likely that many of the RC–APEX bait–prey interactions we identified occurred in the cytoplasm rather than the ring canal. The proximity ligation assay showed ring canal-specific signal for most of the interactions tested, but there were also positive foci present in the cytoplasm. Cytoplasmic interactions could be important for assembling protein complexes destined for ring canals; for example, HtsRC could bind to Filamin in the cytoplasm before localizing to ring canals. We expect that additional analysis of the RC–APEX preys will lead to new insights about ring canal formation, maintenance, and growth during development.

## Materials and Methods

### Drosophila Husbandry

Drosophila were maintained at 25°C on standard fly food medium. Prior to ovary dissections, females were fattened on wet yeast paste overnight at 25°C. See Key Resources Table for a detailed list of fly stocks used in this study.

### Molecular Cloning and Drosophila Transgenesis

#### (BAC) pav::HA::APEX2::FLAG

APEX2 was recombineered into a BAC containing the *pav* gene at its C-terminus (BAC ID 322-102N3) to create *pav::HA::APEX2::FLAG*. This BAC contains the entire *pav* locus on a 21 kb genomic fragment (chr3L:4,229,286…4,250,505, FlyBase release 6). Briefly, we used a 2-step BAC recombineering protocol to first insert a Kanamycin resistance cassette (Wang et al., 2006) which was subsequently replaced by *HA::APEX2::FLAG* through streptomycin selection. The *HA::APEX2::FLAG* sequence was made through PCR amplification of *APEX2* from *pcDNA3–APEX2-NES* (Addgene Plasmid #49386) with primers designed to add an N-terminal *HA* epitope tag sequence and a C-terminal *FLAG* epitope tag sequence. Note that the *APEX2* sequence contained a C-terminal nuclear export signal (NES), which was incorporated in an attempt to keep Pav::APEX localized exclusively to ring canals. The final plasmid was injected into BL#24872 into the attP3B site on chr2L at Rainbow Transgenic Flies, Inc. (Camarillo, CA).

#### otu–ovhts::V5::APEX1

The *V5::APEX1* coding sequence was amplified from *pcDNA3–mito-V5::APEX1* (Rhee et al., 2013; Addgene Plasmid #42607) using primers designed to create a *V5::APEX1* fragment flanked by 3’ XhoI and 5’ NotI restriction sites. *pCOH–ovhts::GFP* (Petrella, Smith-Leiker, and Cooley, 2007) was digested with XhoI and NotI to excise *GFP* and the *V5::APEX1* fragment was ligated into the plasmid in-frame and in place of *GFP*. Transgenic flies were generated via P-element-mediated insertion at GenetiVision Corp. (Houston, TX).

#### UASp–APEX2::V5::kelch

The *APEX2* coding sequence was amplified from *pcDNA3–APEX2-NES* (Addgene Plasmid #49386) and the entire *kelch* coding sequence was amplified with primers designed to contain an N-terminal *V5* tag sequence. Through overlap-extension PCR, the *APEX2::V5::kelch* coding sequence was assembled with flanking attB1 and attB2 sites and recombined first into *pDONR201* and then into *pPW–attB* (described above) in BP Clonase II and LR Clonase II Gateway recombination reactions, respectively. The *pPW–APEX2::V5::kelch* plasmid was injected into a strain carrying the attP2 phiC31 landing site on chr3L at Rainbow Transgenic Flies, Inc. (Camarillo, CA).

#### UASp–APEX2::V5::KREP

The *APEX2* coding sequence was amplified from *pcDNA3–APEX2-NES* (Addgene Plasmid #49386) and the *KREP* coding sequence of *kelch* was amplified with primers designed to contain an N-terminal *V5* tag sequence. Through overlap-extension PCR, the *APEX2::V5::kelch* coding sequence was assembled with flanking attB1 and attB2 sites and recombined first into *pDONR201* and then into *pPW–attB* (described above) in BP Clonase II and LR Clonase II Gateway recombination reactions, respectively. The *pPW–APEX2::V5::kelch* plasmid was injected into a strain carrying the attP2 phiC31 landing site on chr3L at Rainbow Transgenic Flies, Inc. (Camarillo, CA).

#### UASp–GBP::FLAG::APEX2

The *GBP* (Rothbauer et al., 2008) coding sequence was amplified from with primers designed to contain 3’ and 5’ Sfil restriction sites. The *FLAG::APEX2* sequence was amplified from *pcDNA3–APEX2-NES* (Addgene Plasmid #49386) with primers designed to contain a 3’ Sfil site and a 5’ BamHI site. The fragments were digested with their respective restriction enzymes and ligated into a *pUASp-attB* plasmid modified to contain a polylinker sequence downstream of *UASp* site and P-element transposons. Transgenic flies were generated via P-element-mediated insertion at Rainbow Transgenic Flies, Inc. (Camarillo, CA).

### Tissue fixation, fluorescence microscopy, and imaging

See Key Resources Table for more information regarding antibodies and reagents used in this study. Ovaries were dissected in IMADS buffer (ionically matched Drosophila saline; Singleton and Woodruff, 1994) and fixed for 10 minutes in either 6% formaldehyde, 75 mM KCl, 25 mM NaCl, 3 mM MgCl2, and 17 mM potassium phosphate, pH 6.8 (Verheyen and Cooley, 1994) or 4% formaldehyde in PBT (phosphate-buffered saline with 0.3% Triton X-100 and 0.5% BSA). Fixed tissue was washed in PBT and incubated with indicated primary antibodies in PBT for ~1-2 hours at room temperature or overnight at 4°C. Following washes with PBT, tissue was incubated with secondary antibodies conjugated to Alexa Fluors and DAPI in PBT for ~1-2 hours at room temperature. F-actin was labeled with phalloidin conjugated to TRITC. Samples were washed in PBT and mounted on slides in ProLong Diamond (Thermo Fisher). Samples were imaged with a Leica SP8 laser scanning confocal microscope using a 40x 1.3 NA oil-immersion objective lens or a Zeiss Axiovert 200m equipped with either a CARV II spinning disc confocal imager or a CrEST X-light spinning disc system and a Photometrics CoolSNAP HQ2 camera using a 40x 1.2 NA water-immersion objective.

### Western analysis of ovary lysates

Ovary lysates were generated by homogenizing dissected ovaries in SDS-PAGE sample buffer. One ovary equivalent was loaded per lane and separated by SDS-PAGE. Proteins were transferred to nitrocellulose membrane, stained with amido black to visualize total protein, blocked in 5% milk in TBST (Tris-buffered saline with 0.1% Tween-20) for 30 minutes at room temperature, and probed with the indicated primary antibodies for ~2 hours at room temperature in 5% milk/TBST. After washing blots three times in TBST for five minutes, the blots were incubated with HRP-conjugated secondary antibodies for ~1 hour at room temperature followed by five minutes of ECL development. A CCD camera was used for imaging. See Key Resources Table for more information on the antibodies used in this study.

### Proteomic analysis of APEX samples

#### In *vivo* biotinylation of proteins using RC–APEX baits

Prior to dissection, female flies were fattened overnight on wet yeast paste at 25°C in a vial with males. For small-scale pilot experiments or experiments in which fluorescence microscopy was the final experimental readout, ovaries from 10–20 flies were used for biotin-phenol proximity labeling. For large-scale proteomic experiments, three batches of labeling were performed with ovaries from 30-35 flies in each batch, yielding a total of 100 flies used per sample. Ovaries were dissected in 1x PBS, pH 8.0 and gently pipetted up and down ~10 times in order to splay open the ovaries. The pH of all solutions was kept above 7.5 to prevent precipitation of the biotin-phenol. Egg chambers were incubated in 500 *μ*L Biotin-phenol Labeling Solution (1 mM biotin-phenol, 0.5% digitonin, 1x PBS pH 8.0) for three minutes with rotation at room temperature. To initiate APEX-mediated biotinylation, H_2_O_2_ was added to a final concentration of 0.05 mM and the solution was gently inverted to mix for 30 seconds. After 30 seconds, 10 mM NaN_3_ was added to quench the reaction. Egg chambers were washed 3x in 500 *μ*L Quenching Buffer (10 mM NaN_3_ in 1x PBS pH 8.0). For every labeling reaction performed, egg chambers were always fixed and analyzed by fluorescence microscopy to ensure that the labeling reaction was successful (i.e. that ring canals were biotinylated). Standard fixation and fluorescence protocols were used, as described above. To visualize biotin signal in egg chambers, fixed and permeabilized egg chambers were incubated with streptavidin-AF488 or streptavidin-AF568 at a 1:500 dilution in PBT for ~1 hour at room temperature. For large-scale labeling reactions, a very small amount of egg chambers were removed after the last wash in Quenching Buffer and analyzed by fluorescence microscopy. Immediately after the last wash, the remaining Quenching Buffer was removed from the egg chambers, and the tissue was frozen at −80°C.

#### Purification of biotinylated proteins from ovary lysates

Labeled and quenched egg chambers were thawed on ice and the tissue from the 3 batches of labeling was combined into one tube for each sample. Tissue was ground in 100-200 *μ*L Urea Lysis Buffer (8M Urea, 10 mM Tris pH 8.0, 50 mM NaPi, 300 mM NaCl, 5 *μ*g/mL each chymostatin, leupeptin, antipain, pepstatin, 10 mM NaN_3_,1 *μ*M Bortezomib, 10 mM N-ethylmaleimide, 5 mM Trolox, 10 mM Sodium Ascorbate) with a mechanical homogenizer. After grinding, additional Lysis Buffer was added to a final volume of 1 mL and the lysate was incubated for an additional 30 minutes at room temperature to ensure for sufficient lysis and solubilization. The crude lysate was centrifuged at 15,000 rpm for 10 minutes at room temperature, and the clarified lysate was transferred to a new tube with 500 *μ*L pre-equilibrated magnetic streptavidin beads and incubated for 1 hour at room temperature with rotation. The unbound fraction was removed from the beads and the beads were washed three times in 1 mL Urea Wash Buffer (2 M Urea, 10 mM Tris pH 8.0, 50 mM NaP_i_, 300 mM NaCl, 5 *μ*g/mL each chymostatin, leupeptin, antipain, pepstatin, 10 mM NaN_3_,1 *μ*M Bortezomib, 10 mM N-ethylmaleimide, 5 mM Trolox, 10 mM Sodium Ascorbate). In order to analyze the purified fraction by western blotting, 100 *μ*L bead/wash solution was removed during the last wash (i.e. 10% of bead fraction). Beads were eluted with vigorous mixing at 95°C in 50 *μ*L Elution Buffer (SDS sample buffer (0.1 M Tris pH 8.0,4% SDS, 0.01% Bromophenol blue, 5% *β*-mercaptoethanol) with 10 mM biotin). Following the last wash, the remaining wash solution was removed from the rest of the beads and the beads were frozen immediately at −80° C or on dry ice. Beads were shipped on dry ice overnight to MS Bioworks (Ann Arbor, MI) for mass spectrometry analysis using their IP-works platform.

#### Sample preparation for mass spectrometry

Proteins were eluted from streptavidin beads by incubation with 1.5x LDS buffer for 15 minutes at 100°C. Eluted proteins were separated from the beads by centrifugation and magnetic separation. 50% of each eluate was processed by SDS-PAGE with a 10% Bis-Tris NuPAGE gel using the MES buffer system (Invitrogen). The gel lanes were excised, sliced into 10 equal sized fragments, and subjected to in-gel trypsin digestion using a robot (ProGest, DigiLab). For the in-gel trypsin digestion, gel slices were washed with 25 mM ammonium bicarbonate followed by acetonitrile, reduced with 10 mM dithiothreitol at 60°C, alkylated with 50 mM iodoacetamide at room temperature, digested with sequencing grade trypsin (Promega) at 37°C for 4 hours, and quenched with formic acid. The resultant supernatant was analyzed by mass spectrometry.

#### LC-MS/MS data acquisition and processing

Half of each digested sample was analyzed by nano LC-MS/MS with a Waters NanoAcquity HPLC system interfaced to a ThermoFisher Q Exactive. Peptides were loaded on a trapping column and eluated over a 75 *μ*m analytical column at a flow rate of 350 nL/min. Both columns were packed with Luna C18 resin (Phenomenex). The mass spectrometer instrument was operated in data-dependent mode with the Orbitrap operating at 780,000 FWHM for MS and 17,500 FWHM for MS/MS. The fifteen most abundant ions were selected for MS/MS. Data were searched using a local copy of Mascot (Version 2.6.0; Matrix Science) using the UniProt Drosophila melanogaster database with common laboratory contaminants added as well as custom protein sequences for HtsRC that included known SNPs and an N-terminal cleavage site at V693 and APEX1 and APEX2 protein sequences. Trypsin was selected as the enzyme and two missed cleavages were allowed. Cysteine carbamidomethylation was selected as a fixed modification. Methionine oxidation, N-terminal acetylation, N-terminal pyroglutamylation, asparagine or glutamine deamidation, and tyrosine biotinylation by biotin-tyramide (chemical formula: C_8_H_23_N_3_O_3_S) were selected as variable modifications. A peptide mass tolerance of 10 ppm and a fragment mass tolerance of 0.02 Da were used. The resultant Mascot DAT files were parsed using Scaffold (Version 4.8.4; Proteome Software) for validation, filtering, and to create a non-redundant protein list per sample. Experiment-wide grouping with binary peptide-protein weights was selected as the Protein Grouping Strategy. Peptide and Protein Thresholds were set to 95% and 99%, respectively, with a 1 peptide minimum. These settings resulted in a 0.0% Peptide FDR and a 0.4% Protein FDR as measured by Decoy matches. The Scaffold Samples report was exported as an Excel spreadsheet. Contaminant proteins were removed from the spreadsheet dataset. Spectral peptide matches for the Ovhts polyprotein encoded by the *ovhts* transcript were manually parsed apart into its two corresponding protein products: HtsF (AA 1-658) and HtsRC (AA 659-1156). Thus, spectral counts were manually entered into the spreadsheet for HtsF and HtsRC proteins across each sample.

#### Data analysis using SAINT

The Automated Processing of SAINT Templated Layouts (APOSTL) (Kuenzi et al., 2016) tool was used to pre-process the MS data to generate corresponding data files that are compatible with analysis by SAINTexpress (the new-and-improved SAINT tool; see Teo et al., 2014). The resultant SAINTexpress dataset was processed further using APOSTL; namely, normalized spectral abundance factor (NSAF) values were calculated for each prey based on its respective SpC observed in each bait as well as the log_2_ fold-change compared to control (log_2_(FoldChange)).

#### Interactome network visualization with Cytoscape

Cytoscape was used to display the ‘interactomes’ of the RC–APEX baits and their respective high-confidence preys. RC–APEX baits and their prey genes were indicated by colored squares and circles, respectively. The size of the prey gene circle was scaled based on its average abundance compared to control samples (log_2_(FoldChange) NSAF). Known ring canal genes were manually denoted with a solid outline. The Molecular Interaction Search Tool (MIST) (Hu et al., 2018) tool was used to search for previously-established interactions between all high-confidence prey genes, and positive interactions were indicated with a dotted line.

#### Generation of dot plots and heatmaps with ProHits-viz

The ProHits-viz tool (Knight et al., 2017) was used to generate dotplots to visualize proteomics data for any high-confidence prey across all four RC–APEX bait samples. The color of each prey ‘dot’ was dependent on the NSAF log_2_(FoldChange) for each bait compared to the control samples. The size of each dot is dependent on its relative abundance between bait samples, meaning that the maximum-sized dot for each prey is allocated to the largest SpC value within that bait and the other dots are scaled proportionately. High-confidence preys within each bait have a yellow outline, corresponding to a SAINTscore equal to or greater than 0.8, while preys with a SAINTscore below 0.8 for that bait have a faint outline.

#### Generation of volcano plots

Volcano plots were generated for each RC–APEX bait in order to visualize the quantitative proteomic data relative to its statistical significance. For each protein identified by MS, a p-value was calculated for its spectral counts within the biological replicates for each RC–APEX bait and control samples. For all identified proteins, the −log_10_(p-value) was plotted against the NSAF log_2_(FoldChange) compared to controls for each RC–APEX bait sample.

### Validation of RC–APEX high-confidence preys using proximity ligation assay

Ovaries from →50 flies were dissected and fixed in 6% formaldehyde, 75 mM KCl, 25 mM NaCl, 3 mM MgCl2, and 17 mM potassium phosphate, pH 6.8 with heptane for eight minutes. After washing three times for 10 minutes in PBT, the tissue was incubated with 500 *μ*L 30% sucrose solution with gentle rotation for 30 minutes, at which point the ovaries settled to the bottom of the tube. Tissue was rinsed with fresh sucrose solution and transferred to an Eppendorf tube cap where it was fully submerged in cold OCT compound. After a 1-2 hour incubation, the tissue was embedded in a cryoblock mold in OCT by incubating on dry ice for 10 minutes. Cryoblocks were stored at −80°C until sectioning. Prior to sectioning, blocks were equilibrated at −20°C for five minutes. 20 *μ*m slices were sectioned, transferred to Superfrost Plus charged slides (Fisher Scientific), and stored at −80° C until future use. For the PLA, the Duolink^®^ In Situ Red Starter Kit Mouse/Rabbit (Sigma-Aldrich) was used with minor protocol modifications. Slides containing ovary cryosections were equilibrated at room temperature for five minutes, then blocked for one hour in PLA Blocking Buffer (1x PBS, 0.3% Triton X-100,1% BSA). An Aqua Hold II pen was used to form hydrophobic barriers around tissue on the slide to contain the small volumes of solutions used in this protocol. Slides were placed in a humidity chamber and tissue was incubated with 40 *μ*L primary antibody solution and FITC-phalloidin in PLA Blocking Buffer for two hours at room temperature. See Key Resources Table for a list of antibodies and their concentrations used in this assay. Tissue was washed twice for five minutes with PLA Blocking Buffer and incubated in 40 *μ*L PLA Probe Solution (8 *μ*L PLUS probe, 8 *μ*L MINUS probe, 24 *μ*L Antibody Diluent) for 1.5 hours at 37° C in the humidity chamber. Slides were washed twice for five minutes in Wash Buffer A then incubated for 45 minutes in the humidity chamber at 37° C in 40 *μ*L Ligation Solution (8 *μ*L Ligation Buffer, 31 *μ*L ddH_2_O, 1 *μ*L Ligase). Slides were washed twice for five minutes in Wash Buffer A and incubated for two hours in the humidity chamber at 37° C in 40 *μ*L Amplication Solution (8 *μ*L Amplification Buffer, 31.5 *μ*L ddH_2_O,0.5 *μ*L Polymerase) supplemented with FITC-phalloidin. Slides were washed twice for 10 minutes in Wash Buffer B, rinsed for one minute with 0.01x Wash Buffer B, and mounted in 30 *μ*L PLA Mounting Medium overnight in the humidity chamber at 4°C. Slides were imaged on a Leica SP8 laser scanning confocal microscope using a 40x 1.3 NA oil-immersion objective lens and all laser settings were equal between each slide and its respective control to allow for quantitative comparison of PLA foci. Positive PLA foci were counted in FIJI using the Auto-thresholding and Particle Analysis tools, with the minimum area of a foci set to 2 *μ*m^2^. Egg chamber area was calculated in FIJI using the Freehand and Area Measure tools.

### Validation of prey genes by RNAi screening

See Key Resources Table for a list of fly stocks used for the RNAi screen. Ovaries were dissected from fattened females, fixed, and stained with TRITC-phalloidin and anti-HtsRC antibody to check for any abnormalities in egg chamber development or ring canal morphology.

## Acknowledgments

We thank Peter McLean for help with the (*BAC*) *pav::APEX* transgene. We thank Tian Xu (Yale University, Westlake University) for providing access to the Leica SP8 confocal microscope. We are grateful to Ellen LeMosy (Augusta University) as well as members of the Weatherbee lab (Yale University) for help with the cryosectioning protocol. We thank Alice Ting for helpful comments and access to protocols for APEX-mediated biotinylation. We are grateful to Michael Ford and the MS Bioworks team for their services. Stocks obtained from the Bloomington Drosophila Stock Center (NIH P40OD018537) were used in this study. We thank the TRiP at Harvard Medical School for providing transgenic RNAi fly stocks used in this study. K.M.M. was a Cold Spring Harbor Proteomics Course participant. K.M.M. and R.S.K. were supported in part by the National Institute of General Medical Sciences NIH training grant T32 GM007223 and a Gruber Science Fellowship. This work was funded by NIH grants: R01 GM043301 and RC1 GM091791.

## Author Contributions

Conceptualization, K.M.M., L.C.; Investigation, K.M.M., R.M.S., R.S.K., L.C.; Methodology, K.M.M., R.M.S., R.S.K., L.C.; Visualization - K.M.M., R.M.S., L.C.; Funding Acquisition, L.C.; Resources, K.M.M., R.M.S., R.S.K., L.C.; Writing – Original Draft, K.M.M.; Writing – Review & Editing, K.M.M., R.M.S., R.S.K., L.C.; Supervision, L.C.

## Declaration of Interests

These authors have no competing interests.

## Supporting Information

**Figure S1.**
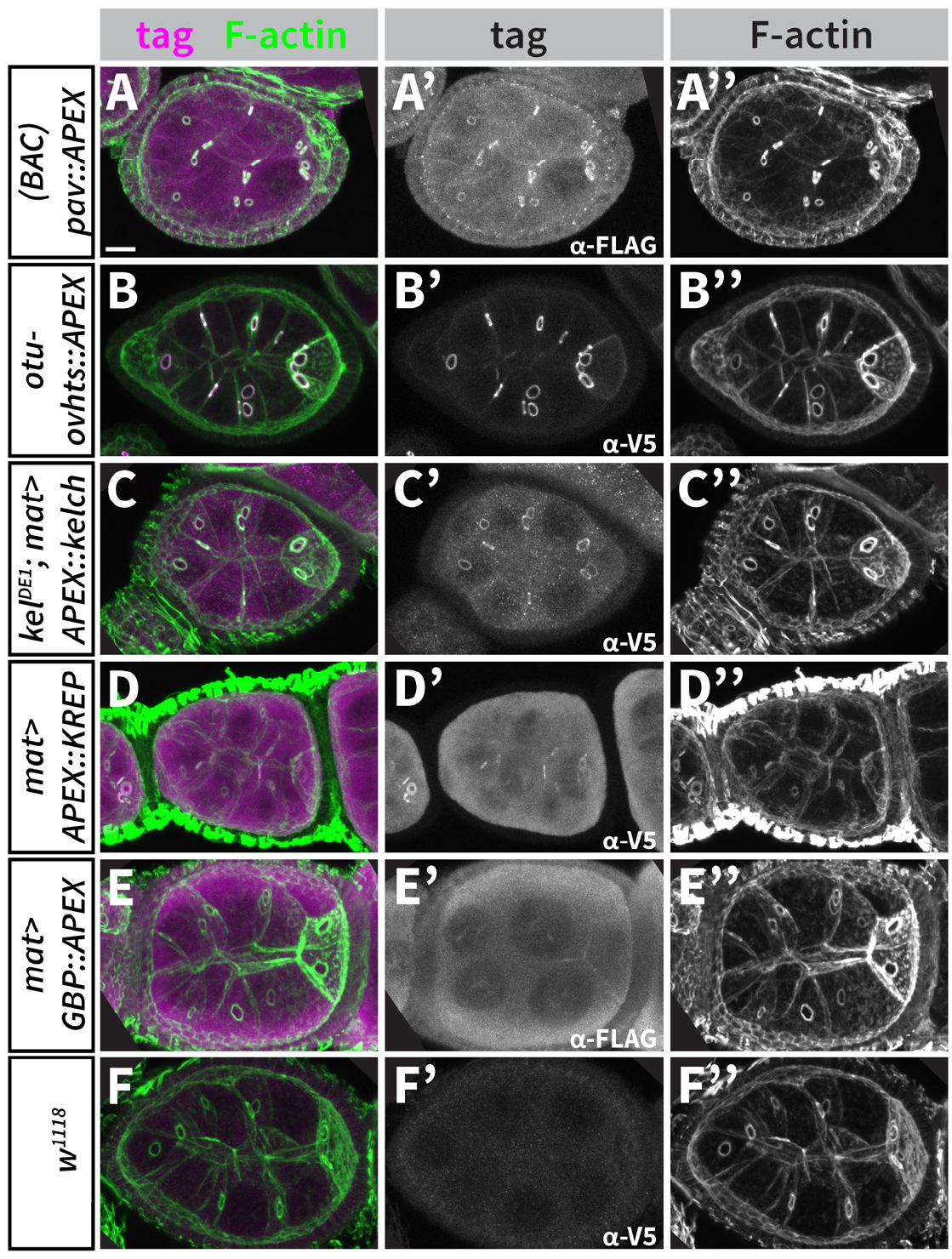
RC–APEX baits localize to ring canals in early-stage egg chambers. (A-F) Stage 6 egg chambers expressing various RC–APEX baits, as indicated by genotypes listed on the left, were stained with TRITC-phalloidin (A”-F”) and their respective epitope tags (A’-F’) to visualize and confirm proper RC–APEX bait construct localization to ring canals. Scale bar, 20 *μ*m.

**Figure S2.**
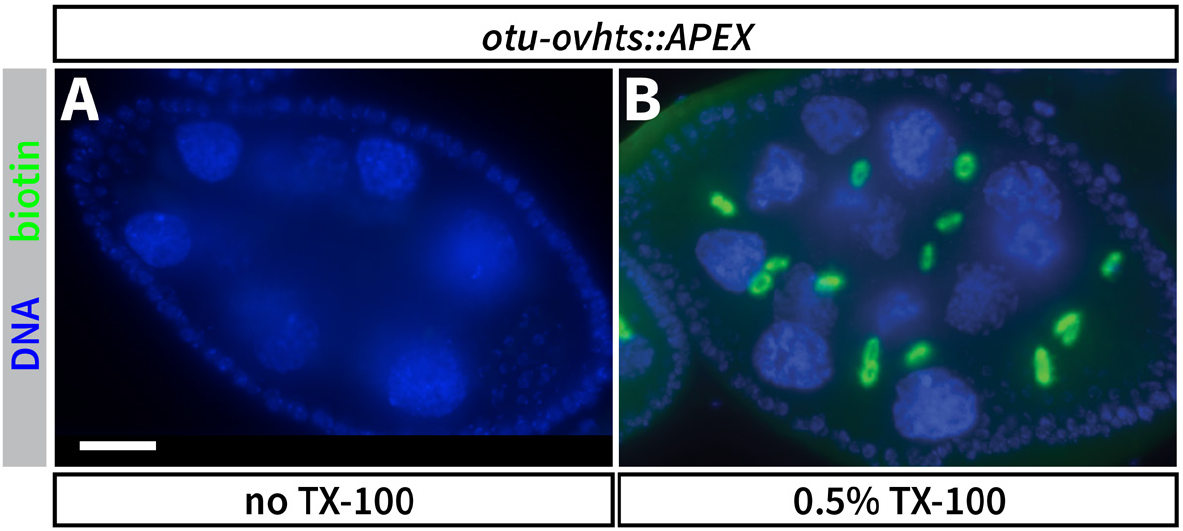
Permeabilization of ovary tissue with detergent is required for biotin labeling. Egg chambers expressing HtsRC::APEX were pre-treated without (A) or with 0.5% TX-100 in PBS pH 8.0 (B) for 10’ at room temperature to permeabilize the cell membranes to allow for entry of the membrane-impermeable biotin-phenol. The labeling reactions were initiated after addition of 500 *μ*m biotin-phenol for 20’, at which point reactions were quenched with 10 mM NaN_3_ and egg chambers were fixed and stained with DAPI and streptavidin-AF488 to reveal extent of biotin labeling. TX-100 treatment allowed for robust APEX-mediated biotinylation of ring canals (B). Also note that APEX-mediated biotinylation was achieved without addition of exogenous H_2_O_2_. Scale bar, 20 *μ*m.

**Figure S3.**
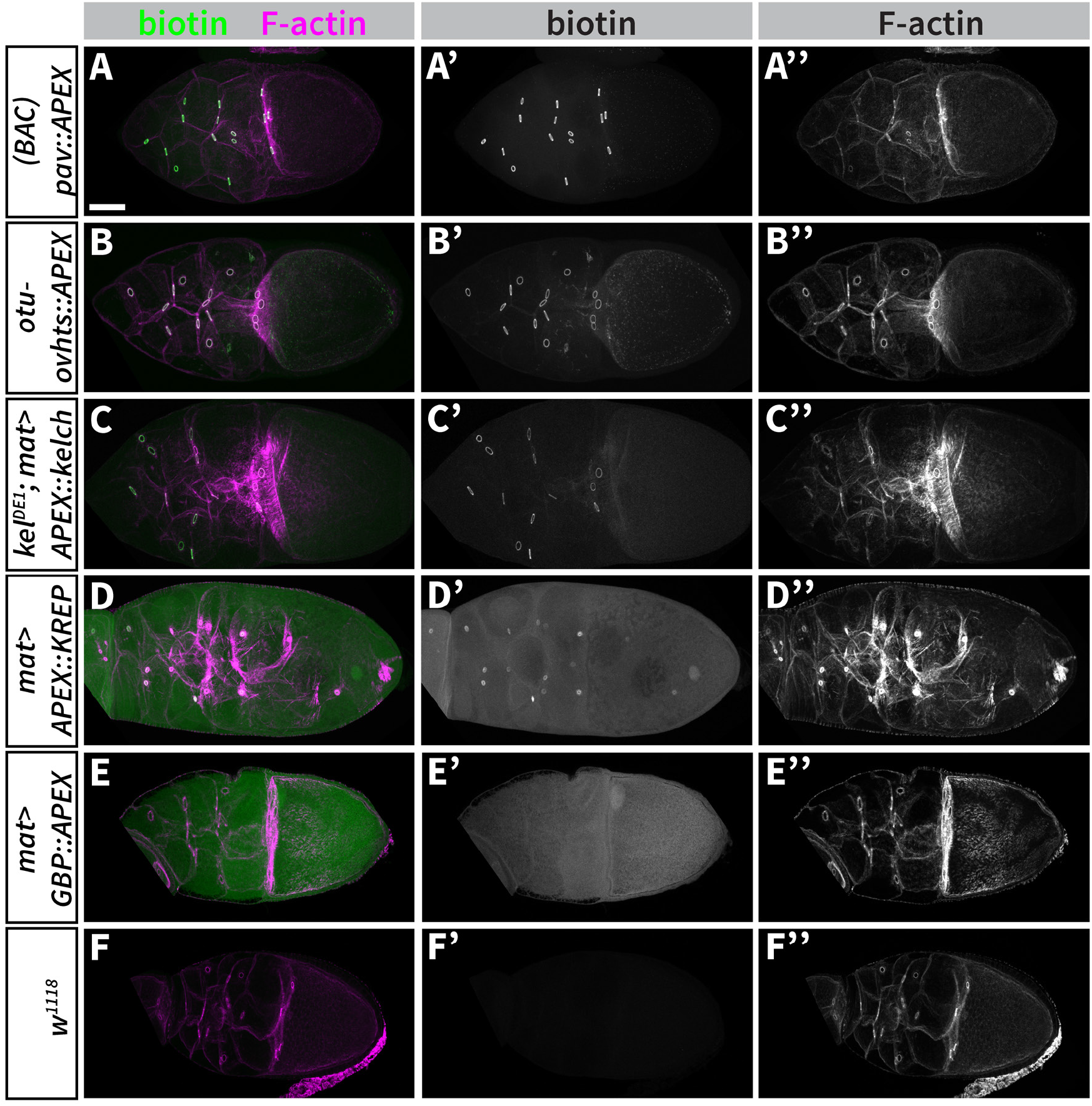
RC–APEX baits specifically biotinylate ring canals in late-stage live egg chambers. (A-F) APEX-mediated biotinylation in live egg chambers was performed as previously described, and egg chambers were fixed and stained with TRITC-phalloidin (A”-F”) and streptavidin-AF488 (A’-F’) to visualize F-actin and biotin. Images are of stage 9 or 10 egg chambers. Scale bar, 50 *μ*m.

**Figure S4.**
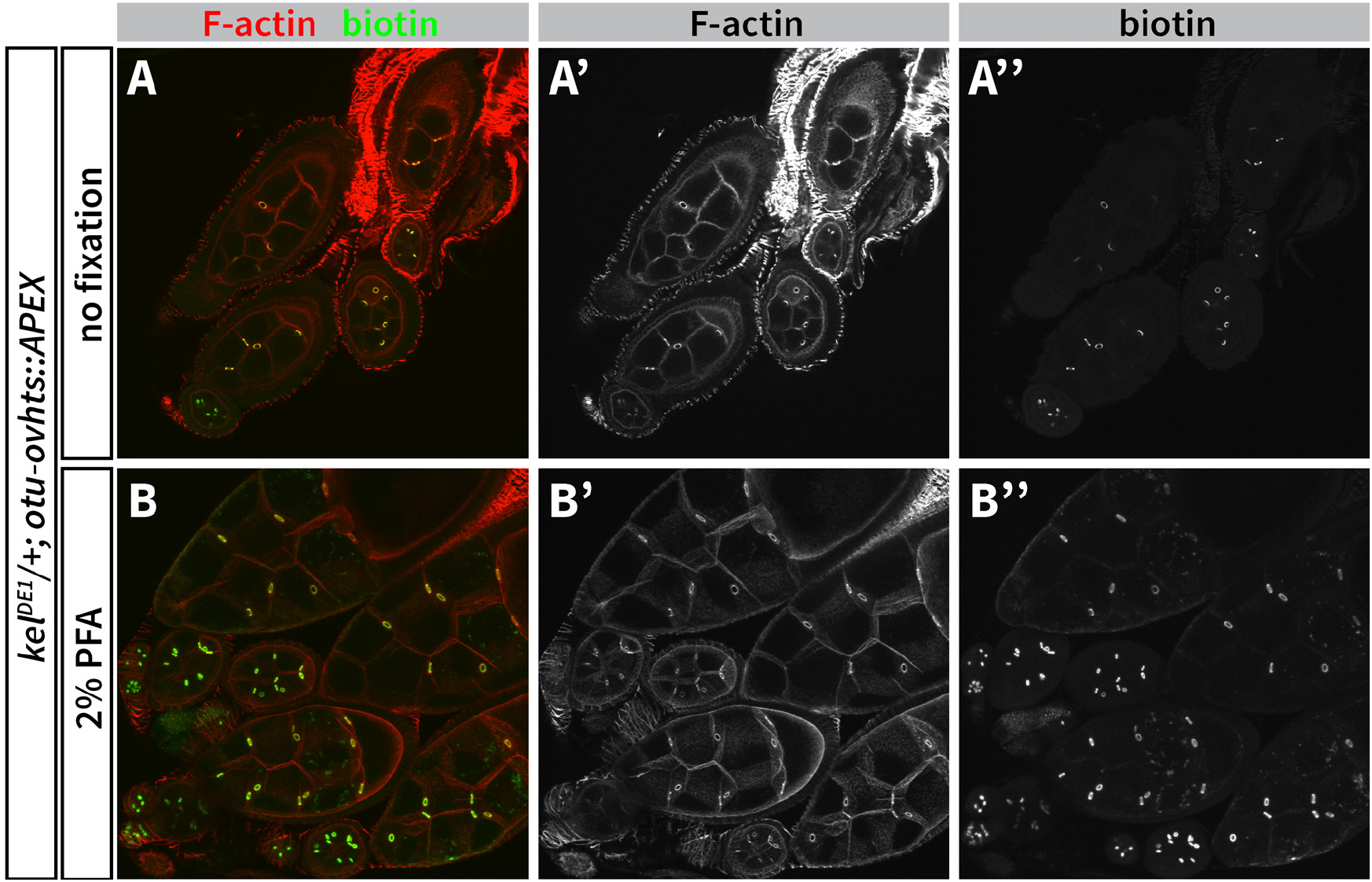
APEX enzyme is active after formaldehyde fixation. (A-B) Ovaries expressing *otu–ovhts::APEX* in a *kel^DE1/+^* background were dissected in PBS pH 8.0 and fixed for 10’ in 2% paraformaldehyde (PFA). After fixation, the tissue was permeabilized by incubating in 0.5% TX-100 for 10’, and APEX-mediated biotinylation was initiated for 20’ after addition of 500 *μ*m biotin-phenol in PBS. Reactions were quenched by 10 mM NaN_3_, and the tissue was again fixed in 4% PFA and stained with TRITC-phalloidin (A’-B’) and streptavidin-AF488 (A”-B”) to visualize egg chamber morphology and extent of biotinylation. Note the presence of biotinylated ring canals (B, B”) even after egg chambers were fixed prior to biotinylation initiation.

**Figure S5.**
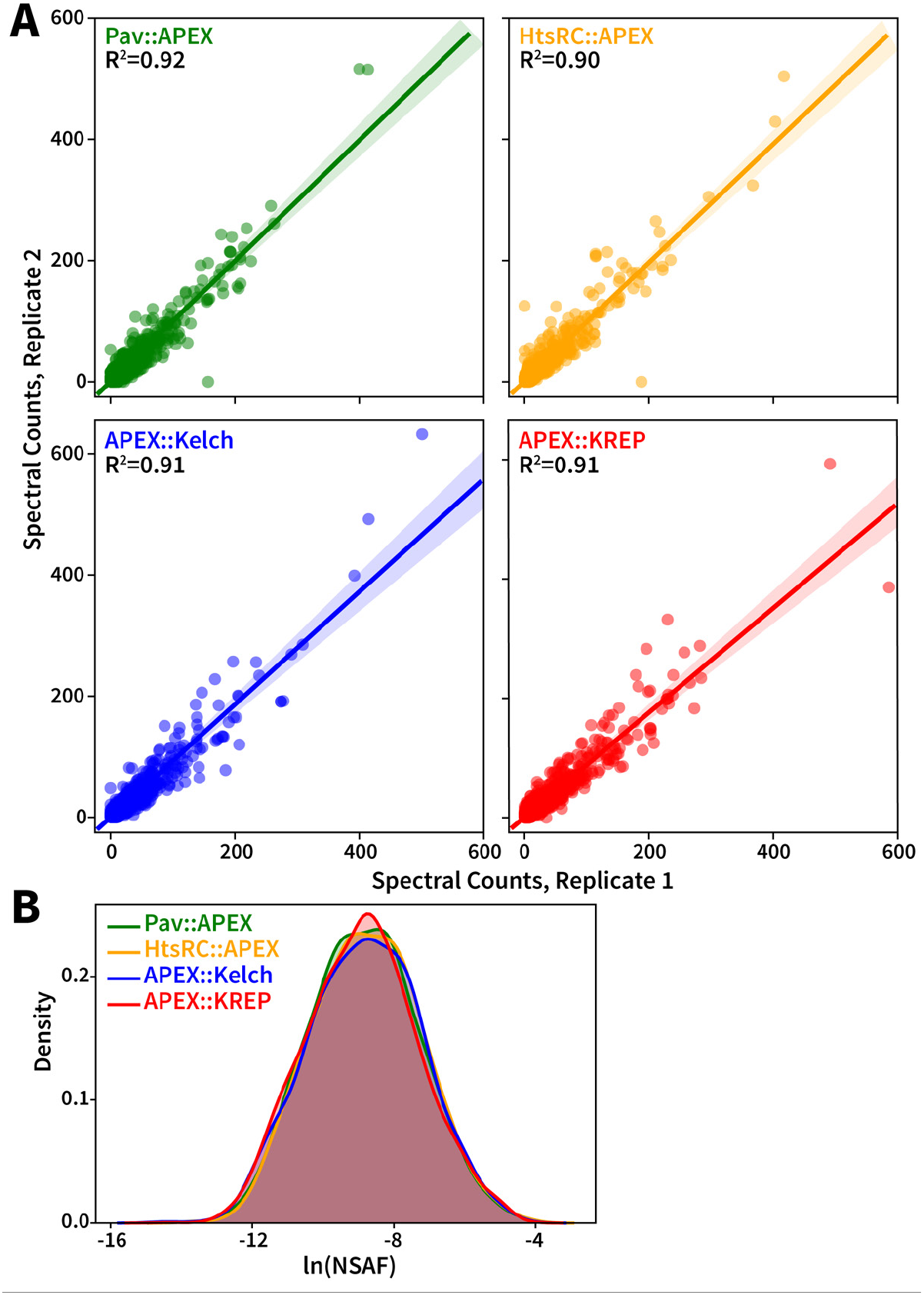
Spectral counts can be used as a semi-quantitative measure of protein abundance for replicate experiments. (A) Spectral counts for each protein identified were plotted for two replicate experiments for all four RC–APEX baits. The solid line and shaded area correspond to the best-fit line and its confidence interval, respectively, for the linear regression analysis, and R^2^ values are indicated on the plots. (B) The frequency of natural log (ln) of normalized spectral abundance factor (NSAF) values of all proteins identified by MS were plotted for all four RC–APEX baits. Note that all of the density plots overlap closely, indicating similar extent of coverage of spectral counts for protein quantitation between the four RC–APEX samples.

**Figure S6.**
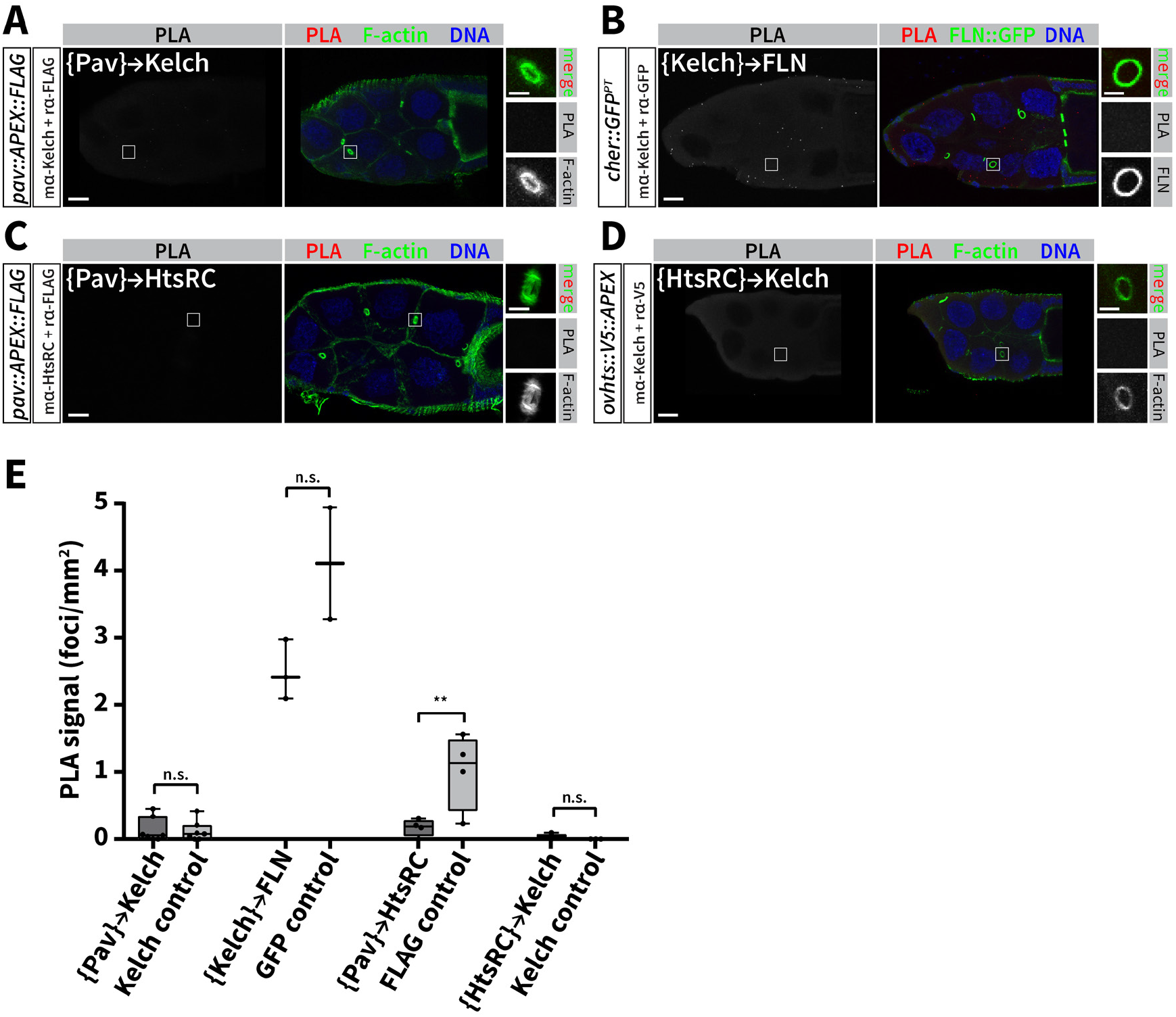
Proximity ligation assay confirms absence of protein-protein interactions between disparate RC–APEX bait protein pairs. (A-D) Proximity ligation assay was performed as described above to test the interaction of RC–APEX bait proteins (Pav, Kelch, and HtsRC) with RC proteins (Pav, Kelch, HtsRC, and Filamin) that were not respective preys and therefore are not expected to be close-proximity interactors. PLA reaction fluorescence was widely absent from egg chambers when testing for interactions between {Pav}→Kelch (A), {Kelch}→Filamin (B), {Pav}→HtsRC (C), and {HtsRC}→Kelch (D), consistent with our proteomics data. (E) FIJI tresholding and particle analysis were used to quantify PLA fluorescent foci in each egg chamber, and foci count is plotted as a function of egg chamber unit area. All interactions tested were negative, meaning there was no enrichment of PLA foci in experimental samples compared to negative controls in which one antibody was omitted.

